# Disrupted astrocyte-endothelial crosstalk drives hemangioblastoma lesions in VHL disease

**DOI:** 10.1101/2025.09.15.675862

**Authors:** Macarena de Andrés-Laguillo, Irene Garcia-Gonzalez, Susana F. Rocha, Aroa Garcia-Cabero, Sandra Ruiz-García, Luis Diago-Domingo, Aimane Danana, Lorena Cussó, Katrien De Bock, José A Enríquez, Rui Benedito

## Abstract

Hemangioblastomas (HBs) are highly vascularized central nervous system (CNS) tumours that can become life-threatening, especially in the context of Von Hippel-Lindau (VHL) disease, caused by the loss of VHL function. The limited pharmacological options targeting VHL-HBs stem from an incomplete understanding of their cellular origin, development, and molecular pathogenesis.

Here we use advanced mouse genetics to show that mosaic deletion of *Vhl* in *Apln*+ cells leads to the formation of precursor tumour-like lesions, composed by clusters of *Vhl-knockout (Vhl^KO^)* astrocytes and surrounding *Vhl-*wild-type *(Vhl^WT^)* vessels that become malformed, resembling early-stage HBs linked to VHL disease. *Vhl^KO^* astrocytes morphologically and transcriptomically resembled the reactive astrocytes characteristic of ischemic CNS injury. They exhibited metabolic rewirement towards glycolysis and upregulation of cell growth pathways. They also expressed several secreted proangiogenic molecules that activate and prevent the normal maturation of neighbouring vessels, leading to VHL-HBs. Temporal conditional genetic analysis revealed that *Vhl* loss need to happen during postnatal development for HBs to form, and that lesions become quiescent in early adulthood. HIF-2α deletion, or MTORC1 inhibition with rapamycin, efficiently inhibited VHL-HBs growth and the associated vascular malformations.

Our work shows that the loss of *Vhl* in single astrocytes induces their growth and pathogenic crosstalk with neighbouring endothelial cells, driving hemangioblastoma development in VHL disease. Our new somatic mosaic mouse models will also enable testing of novel drugs against this disease.

## Introduction

Hemangioblastomas (HBs) are highly vascularized central nervous system (CNS) tumours that, although benign, can become life-threatening due to haemorrhage and the compression effect over the neighbouring neural tissue. HBs account for up to 2.5% of all CNS tumours and can occur sporadically or in the context of von Hippel–Lindau (VHL) disease, in which they are the most common manifestation^1^.

Therapeutic options for the management of HBs are predominantly surgical. While resection is effective, the recurrence of these tumours (particularly in VHL disease) leads to frequent surgical interventions, reducing patients’ life expectancy. Pharmacological therapies remain limited, in part due to an incomplete understanding of the HBs cellular origin, development and molecular pathogenesis^2–4^.

In VHL disease, patients inherit a germline loss-of-function mutation in the hypoxia response repressor gene VHL. Tumour formation requires a somatic second-hit mutation in the remaining wild-type allele. Notably, somatic VHL mutations are also found in ∼78% of sporadic HBs, underscoring the central role of VHL inactivation in both HB forms^5^.

Histologically, HBs are composed of highly abnormal blood vessels that surround the VHL-mutant tumoral mass. However, the vascular component is formed by wild-type VHL+ cells ^5,6^. Interestingly, the origins and identity of the VHL-mutant tumoral cells remains unresolved. While some propose an hemangioblast origin, others have suggested glial cells as potential drivers^7–10^. Although a few animal models have recapitulated some characteristics of HB-like lesions^11,12^, none has studied one of the disease’s most distinctive hallmarks: it’s somatic mosaic nature. Another unresolved question concerns the timing of VHL loss, and the cellular and molecular mechanisms that subsequently induce HB formation. It is unclear whether de novo lesions arise after *VHL* somatic mutations in adulthood, or whether early VHL inactivation gives rise to precursor lesions that later reactivate ^7,13^. Understanding the temporal dynamics of VHL loss and the mechanisms of HB development could help the development of strategies to reduce the progression of VHL disease.

Here, we developed diverse mosaic *Vhl*-knock-out mouse models that recapitulate key histopathological features of VHL-HBs which allowed us to dissect its origins and spatiotemporal mechanisms of development. We identify mutant postnatal astrocytes as drivers of VHL-HB-like lesions. After the loss of VHL in postnatal astrocytes, these become genetically hypoxic and activated, driving not only their own growth but also the abnormalization of the neighbouring blood vessels. The disease development however requires some cellular plasticity, as the mosaic induction of the *Vhl* mutation in adult quiescent organs does not result in HBs.

This work establishes new mouse models and uncovers molecular and cellular mechanisms of VHL disease progression that are relevant for understanding and treating this condition.

## Results

### VHL loss in *Apln-*expressing cells induces Hemangioblastoma lesions

To study the implications of mosaic *Vhl* deletion in HB development, we tested whether the mosaic deletion of *Vhl* in endothelial and hemogenic progenitor cells expressing the *Apln* gene would recapitulate similar pathological features. To this end, we used *iFlpMosaic* mice, which allow us to reliably induce gene knock-out (KO) mosaics without the confounding false positives and false negatives associated with Cre-dependent mosaic genetics^14^. By generating mice containing *Apln-FlpO*^15^*, Tg-iFlp^MTomato-2A-H2B-GFP-2A-Cre/MYFP-2A-H2B-Cherry^* ^14^ and the *Vhl* gene flanked by loxP sites^16^ (here on referred to as *Apln-iFlpMosaic Vhl^flox/flox^*) we could induce ratiometric labelling of both *Vhl^WT^* (MYFP-2A-H2B-Cherry+) and *Vhl^KO^* (MbTomato-2A-H2B-EGFP-2A-Cre+) cells in all *Apln*-expressing cell lineages (**Fig. 1a**). This mouse model represents a unique genetic strategy for inducing *Vhl^KO^* mosaics, considering that *Vhl* is very close and genetically linked to the Rosa26 locus, which so far has prevented its combination with standard Rosa26 knock-in reporter alleles and related lineage-tracing methods.

**Figure 1:**
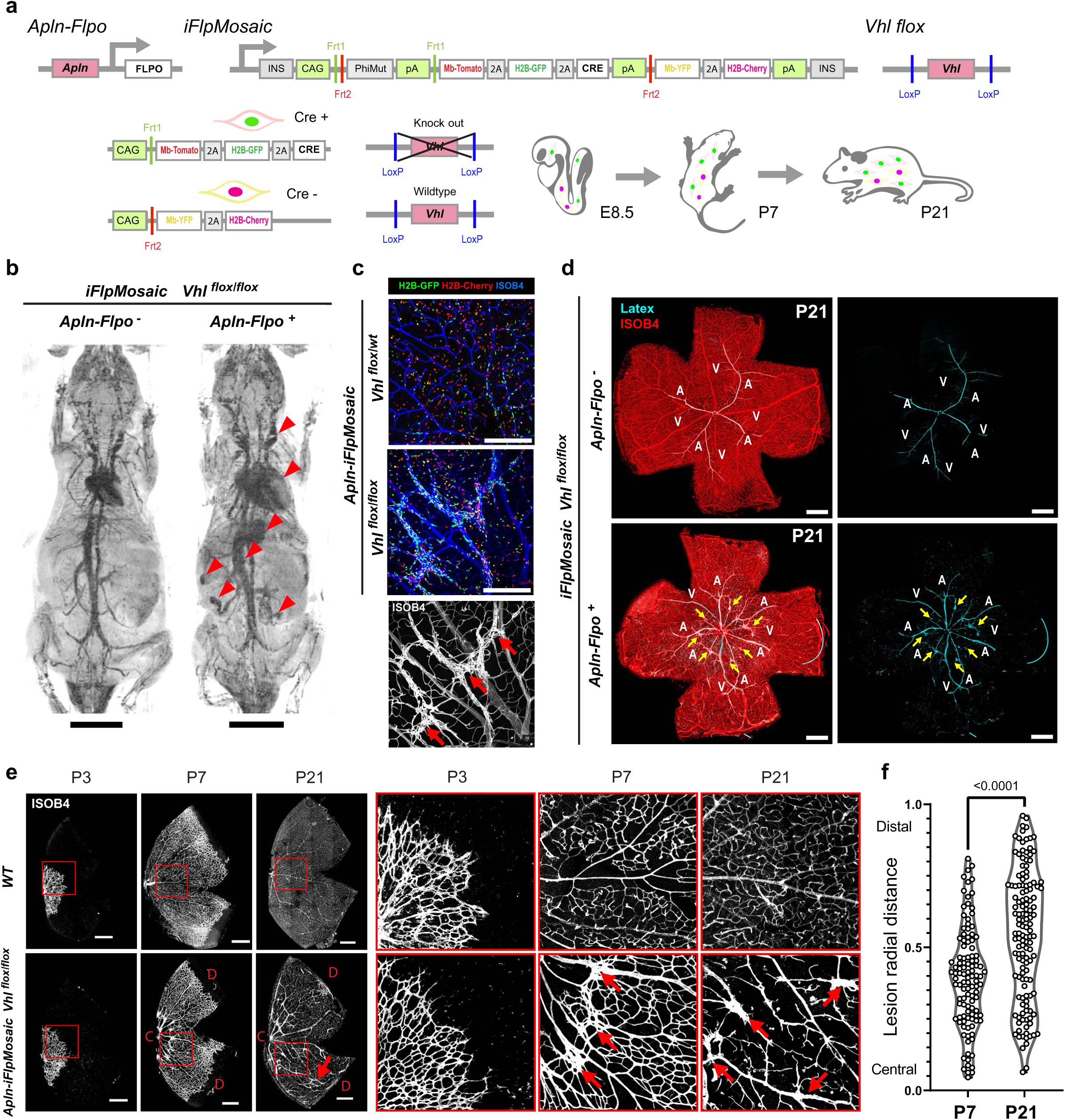
*Apln* driven mosaic deletion of *Vhl* drives Hemangioblastoma lesions. **a,** Tg-*iFlp^MTomato-2A-H2B-GFP-2A-Cre/MYFP-2A-H2B-Cherry^* mosaics were induced with *Apln-FlpO* (recombines endothelial progenitors and derived hematopoietic cells at E8.5) on control heterozygous (*Vhl^flox/wt^*) or homozygous (*Vhl^flox/flox^*) backgrounds to mimic human somatic *Vhl* deletion. **b,** Micro-Computed Tomography vascular imaging shows the macrovasculature. Red arrows indicate vascular alterations in *Apln-iFlpMosaic Vhl^flox/flox^* mice. **c,** Representative confocal micrographs of control and malformed intervein retina areas from 3-week-old *Apln-iFlpMosaic Vhl^flox/wt^*and *Vhl^flox/flox^* mice respectively, immunostained for ISOB4, endogenous GFP (mutant cells) and Cherry (wildtype cells) signal. Red arrows indicate VMs. **d,** Representative confocal micrographs of latex-perfused retinas from the indicated genotypes immunostained for ISOB4 and latex auto-fluorescent detection. Yellow arrows indicate shunts. **e,** Representative confocal micrographs of WT and *Apln-iFlpMosaic Vhl^flox/flox^* retinas immunostained for ISOB4 showing vascular progression and changes in VMs distribution through postnatal days 3, 7 and 21. Red arrows indicate VMs. **f,** Violin plot indicating relative distance of lesions to the center of the retina, each group shows VMs from 4 independent mice. For statistics see Source Data File 1. Scale bars are: b-1cm; c-250μm; d,e-500μm. Abbreviations: A – Artery, V – Vein, C – Central, D – Distal, P3, 7, 21 – Postnatal day 3, 7, 21.

*Apln-iFlpMosaic Vhl^flox/flox^* mice showed increased lethality, with an average life expectancy of 9 weeks (**Extended Data Fig. 1a**). Micro-computed tomography revealed clear cardiomegaly, vessel dilations and angioma-like tortuous vascular structures in adult mutant mice compared to their control littermates (**Fig. 1b**).

Since 60% of VHL patients develop retinal HBs^2,3^, we analysed the mouse retina vasculature of *Apln-iFlpMosaic Vhl^flox/flox^* postnatal day (P) 21 mice. We observed structures that resembled early-stage HBs^17,18^ formed by *Vhl^KO^* (H2B-GFP+) cell clusters and malformed and enlarged blood vessels (**Fig. 1c**). To determine if these vascular malformations (VMs) were perfused, or formed direct shunts between arteries and veins, we performed left ventricle latex perfusion. Given its large particle size and viscosity, latex can only perfuse arteries and not the finely branched capillary plexus downstream, or veins ^19^. However, in the *Apln-iFlpMosaic Vhl^flox/flox^* retinas, the VMs and veins were perfused by latex (**Fig. 1d**), confirming that mosaic loss of VHL cause shunts between arteries and veins.

Besides pathology in the retina, VHL patients frequently manifest cerebellar HBs^2,20^. Using lectin perfusion combined with whole brain imaging after clearing, we also detected malformed vascular structures in the cerebellum from *Apln-iFlpMosaic Vhl^flox/flox^* mice. These structures colocalized with the presence of H2B-GFP *Vhl^KO^* cell clusters (**Extended Data Fig. 1b**). Clearing and imaging of other latex perfused organs from *Apln-iFlpMosaic Vhl^flox/flox^* mice, revealed multisystemic arteriovenous shunts in the kidney, liver and mesentery, which are also reported in human VHL disease ^21^ (**Extended Data Fig. 1c**).

To determine how these lesions develop over time in *Apln-iFlpMosaic Vhl^flox/flox^* mice, we analyzed retinas at P3, P7 and P21. Interestingly, despite mosaic mice having *Vhl^KO^* cells since E8.5^15^, lesions were observed only at P7 and P21 in the retina, but not at P3 (**Fig. 1e**). At P7, lesions were located predominantly in the retina centre, near the optic nerve, where vascular maturation starts and VEGF levels are lower, while they were absent at the distal VEGF-rich angiogenic front. However, at P21, when the whole vascular plexus is mature and quiescent, lesions could be identified also in distal areas, and the distribution was more homogeneous (**Fig. 1f**). Overall, this data suggests that *Vhl^KO^*HB-like lesions are formed during postnatal vascular maturation, in areas with relatively lower VEGF levels, and not during active angiogenesis.

### Hemangioblastomas are caused by the mosaic loss of VHL in non-endothelial cells

To determine the mechanisms driving lesions’ development at high cellular resolution, we analyzed the distribution of the nuclear reporters H2B-GFP (*Vhl^KO^* cells) and H2B-Cherry (*Vhl^WT^* cells). This revealed the presence of large H2B-GFP+ cell clusters within the lesions of *Apln-iFlpMosaic Vhl^flox/flox^* mutant mice (**Fig. 2a,b**). Surprisingly, despite of *Apln* being mostly expressed by ECs (**Extended Data Fig. 2a, b**), most of the *Apln*-derived *Vhl^KO^* cells were negative for the endothelial-specific transcription factor ERG, while *Vhl^Het^* cells from control mosaics were mostly endothelial, as expected (**Fig. 2c**). We took advantage of the ratiometric nature of *iFlpMosaics*^14^ to analyse the ratio between H2B-GFP+ (*Vhl^Het^ or Vhl^KO^*) and H2B-Cherry+ (*Vhl^WT^*) cells, within endothelial (ERG+) and non-endothelial (ERG-) cells. This ratiometric analysis enabled us to spatially quantify how mutant/GFP+ cells expand in relation to neighbouring wildtype/Cherry+ cells. This analysis revealed that, within lesions, H2B-GFP+/*Vhl^KO^* non-ECs were 32 times more frequent than adjacent H2B-Cherry+ (*Vhl^WT^*) non-ECs (**Fig. 2d**). Our control H2B-GFP+/*Vhl^Het^* non-ECs were just 2 times more frequent than H2B-Cherry+ (*Vhl^WT^*) non-ECs, which is expected given the distances between loxP sites (**Fig. 1a**). This data suggests that *Vhl* deletion leads to the expansion of *Apln*-derived non-ECs in the malformed areas.

**Figure 2:**
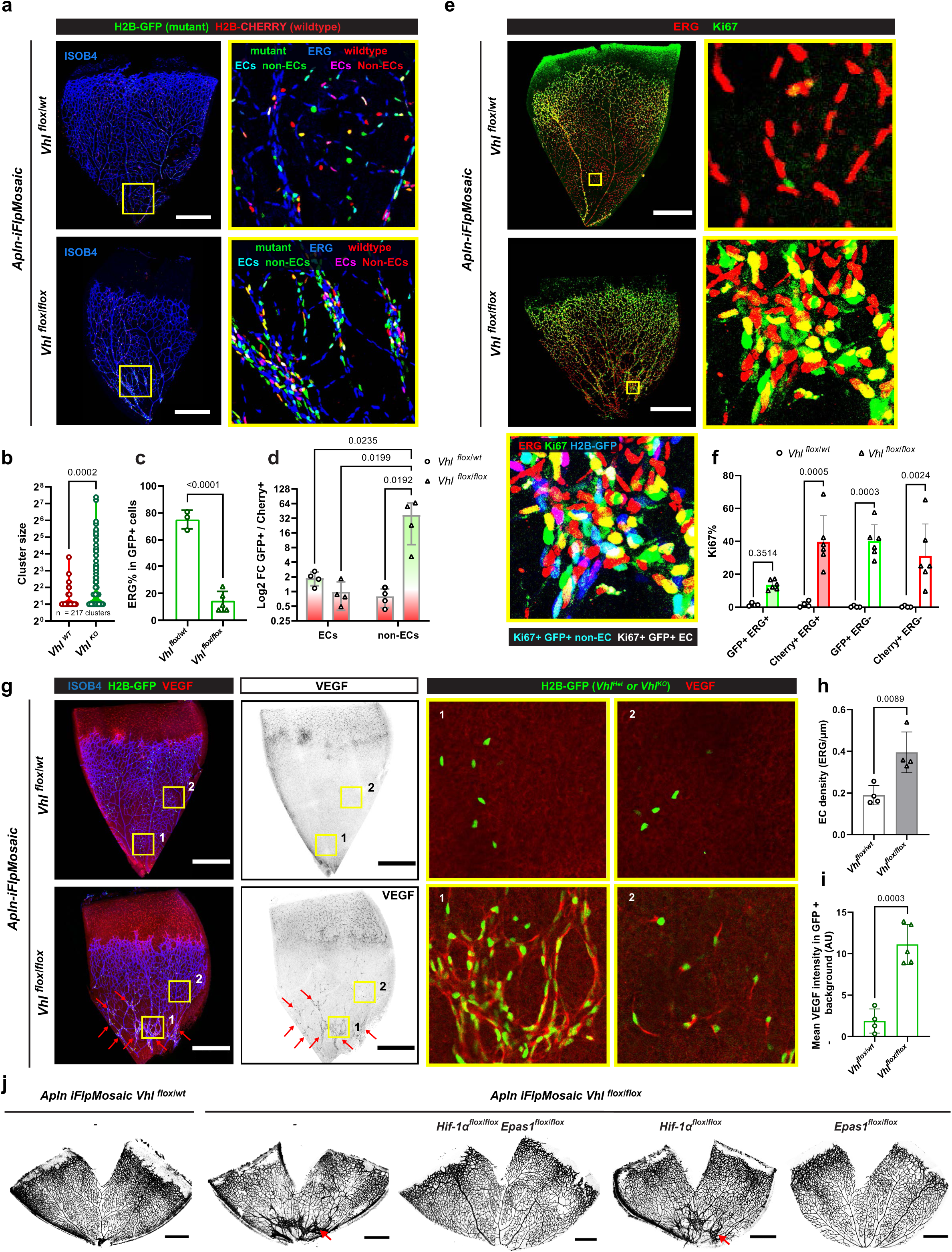
*Apln-iFlpMosaic* HB lesions are formed by highly proliferative *Vhl^KO^* non-EC clusters and malformed angiogenic blood vessels. **a,** Representative confocal micrographs of *Apln-iFlpMosaic* control *Vhl^flox/wt^* and malformed *Vhl^flox/flox^* retinas immunostained for ISOB4, ERG, endogenous GFP (mutant cells) and Cherry (wildtype cells). Magnified insets of central areas show a high frequency of non-EC *Vhl^KO^* cells within HB. **b,** Violin plot indicating cluster size distribution of *Vhl^WT^* (H2B-Cherry) *and Vhl^KO^* (H2B-GFP) cells within *Apln-iFlpMosaic Vhl^flox/flox^* retinas, each group shows 217 clusters from 4 independent mice. **c,** Barplot indicating the percentage of endothelial (ERG+) cells within the GFP+ recombined cells in *Apln-iFlpMosaic* control *Vhl^flox/wt^* vs malformed *Vhl^flox/flox^* retinas. **d,** Barplot displaying the log2 fold change (mutant H2B-GFP vs. WT H2B-Cherry) in *Vhl^flox/wt^* central areas vs *Vhl^flox/flox^*HB-like lesions showing a strong enrichment of *Vhl^KO^* non-ECs in HB-like lesions. **e,** Representative confocal micrographs of *Apln-iFlpMosaic* control *Vhl^flox/wt^* and malformed *Vhl^flox/flox^*retinas immunostained for Ki67, ERG and endogenous GFP (mutant cells). Magnified insets of central areas shown high Ki67 expression in HB-like lesions. **f,** Barplot indicating the percentage of proliferative (Ki67+) in the specified populations within *Vhl^flox/wt^* central areas vs *Vhl^flox/flox^*HB-like lesions. **g,** Representative confocal micrographs of *Apln-iFlpMosaic* control *Vhl^flox/wt^* and malformed *Vhl^flox/flox^* retinas immunostained for ISOB4, VEGF and endogenous GFP signal. Magnified insets of central (1) and distal (2) areas show cell autonomous high expression of VEGF in *Vhl^KO^*cells. **h,** Barplot indicating the endothelial density (ERG+ in vascularized area) in *Vhl^flox/wt^* central areas vs *Vhl^flox/flox^ HB-like lesions*. **i,** Barplot indicating the mean VEGF intensity within GFP+ cells from *Apln-iFlpMosaic* control *Vhl^flox/wt^* and malformed *Vhl^flox/flox^* retinas normalized by background signal. **j,** Representative confocal micrographs of *Apln-iFlpMosaic* retinas with the indicated floxed alleles immunostained for ISOB4, showing that *Epas1*, but not *Hif1a* codeletion prevents HB-like lesions in the *Vhl^flox/flox^* background. Data are presented as mean values +/− SD. For statistics see Source Data File 1. Scale bars are 500μm.

The elevated numbers of non-endothelial H2B-GFP+ (*Vhl^KO^*) cells arising from the Apln-lineage cells within these highly vascularized lesions was unexpected. To dismiss the possibility of Apln-lineage *Vhl^KO^* ECs transdifferentiating into non-ECs (**Extended Data Fig. 2c**), we used the endothelial-specific *Cdh5-CreERT2* line to determine if EC deletion of *Vhl* could lead to labelled non-ECs and HB-like lesions (**Extended Data Fig. 2d**). Postnatal induction of *Cdh5-CreERT2 iSuRe-Cre Vhl^flox/flox^* mice did not cause any transdifferentiaton of ECs or HB-like lesions (**Extended Data Fig. 2e**). Paradoxically, endothelial *Vhl* deletion even delayed angiogenesis and decreased by threefold the frequency of Ki67+ proliferating ECs (**Extended Data Fig. 2f-h)**. Therefore, *Vhl* deletion in ECs reduces their proliferation, whereas its deletion in non-ECs results in their expansion in areas associated to the development of VMs (**Fig. 2a-f**).

As mentioned above, *Apln* is expressed predominantly in ECs but not exclusively. Analysis of postnatal retina scRNAseq data^22^, revealed that several other cell types also express *Apln* at lower levels, such as pericytes, astrocytes or photoreceptors (**Extended Data Fig. 2a,b**). We hypothesized that some of these cell types could increase their numbers due to proliferation upon *Vhl* deletion, accounting for the formation of the non-endothelial *Vhl^KO^* clusters observed, which could then induce VMs by paracrine modulation of the neighbouring ECs (**Extended Data Fig. 2c**). Indeed, both non-endothelial *Vhl^KO^* cells (ERG-negative, H2B-GFP+) and neighbouring wildtype ECs (ERG-positive, H2B-Cherry+), were highly proliferative (40% KI67+) in the lesion areas (**Fig. 2e, f**). This data shows that mosaic deletion of *Vhl* in non-ECs induces their proliferation and stimulates proliferation of the surrounding *Vhl^WT^* vasculature, leading to the formation of VMs, whereas deletion of *Vhl* in ECs themselves inhibits their proliferation (**Extended Data Fig.2h and Fig.2f).**

To elucidate the non-endothelial H2B-GFP+/*Vhl^KO^* signals that could cause an increase in EC proliferation leading to VMs, we first analysed the expression of VEGF, a very strong hypoxia inducible proangiogenic factor that promotes EC proliferation and sprouting^23,24^. *Vhl* loss is expected to increase HIFs stabilization and VEGF expression^24,25^. Immunostaining revealed high levels of VEGF within the H2B-GFP+/*Vhl^KO^* cells from *Vhl^flox/flox^* retinas, which contrasted with the almost undetectable VEGF levels observed in the control *Vhl^flox/wt^* retinas (**Fig.2 g-i).** Individual *Vhl^KO^* cells located in the more distal and angiogenic areas of the retina, where clear VMs were still not present, also showed high VEGF levels (**Fig. 2g right panel).** This confirms that the high production of VEGF by *Vhl^KO^* non-ECs is single-cell autonomous. However, VMs only formed in more mature vascular areas with neighbouring clusters of VEGF-producing *Vhl^KO^* cells.

Altogether, these results indicate that *Vhl* mosaic deletion in non-ECs promotes their entry in cell-cycle (KI67+), triggering expansion and formation of clusters, which then result in higher VEGF levels in the microenvironment surrounding wildtype maturing vessels. The higher VEGF levels likely prevent the normal vascular maturation and remodelling process, with ECs remaining activated and proliferative, leading to enlarged and malformed vessels. These non-EC clusters surrounded by malformed vessels closely resemble the tumoral and vascular component of VHL disease HBs ^5,17,26,27^.

### *Epas1* deletion in *Vhl^KO^* cells prevents the development of hemangioblastomas

Given the observed high VEGF levels present around *Vhl^KO^*cell clusters, we wondered if the development of HB-like lesions could be prevented by pharmacological targeting of VEGF. To test this, we treated *Apln-iFlpMosaic Vhl^flox/flox^* postnatal mice with the VEGF-Trap (Aflibercept/Zaltrap, 25mg/kg) (Saishin et al., 2003) from P3 to P6 (**Extended Data Fig. 3a**). Anti-VEGF treatment reduced, but did not fully prevent, endothelial proliferation in the lesions (**Extended Data Fig. 3b,c**). As a result, there was a tendency to decreased VMs density (**Extended Data Fig. 3d**). Given these results, we hypothesized that a systemic pharmacological VEGF targeting strategy might not be enough to fully block the high local VEGF levels present in the retina, which leads to an incomplete treatment effect. Effective VEGF signalling blocking in the retina can only be achieved by intraocular administration in both humans^2^ and postnatal mice^28^.

To try to prevent lesion development and completely block VEGF production by the genetically hypoxic cells, we instead co-deleted the hypoxia-inducible factors (HIFs). HIF-1α *and* HIF-2α (encoded by *Hif1a* and *Epas1* respectively) are stabilized in the absence of VHL and drive the activation of the hypoxia transcriptional program and VEGF production^25,29^. As expected, at P7, retinas from *Apln-iFlpMosaic Vhl^flox/flox^*, *Hif1a^flox/flox^, Epas1^flox/flox^* mice showed no *Vhl^KO^* clusters or VMs, indicating the essential role of HIFs in HBs development. Interestingly, after deleting each *Hif* gene independently, we observed that only loss of *Epas1*, but not *Hif1a*, rescued the phenotype caused by the *Vhl* loss (**Fig.2j).** The increase in KI67 frequency (**Extended Data Fig. 4a**), *Vhl^KO^*(GFP) cell clustering (**Extended Data Fig. 4b,c**) and VEGF expression (**Extended Data Fig. 4d,e**) was fully prevented by *Epas1* deletion, but not *Hif1a* deletion.

These results indicate that *Epas1*, but not *Hif1a*, is essential for HB-like lesions’ development after *Vhl* loss. Interestingly, clinical trials using the EPAS1/HIF-2A inhibitor Belzutifan are showing great efficiency in the treatment of VHL-HBs^4,30,31^.

### Astrocyte-specific postnatal deletion of *Vhl* induces hemangioblastomas

Given that most *Vhl^KO^* cells in the *Apln-iFlpMosaic* model were not ECs, we next wanted to determine their identity. We first assessed which cells in the postnatal retina express high levels of *Epas1*. For this, we analysed publicly available scRNAseq data^22^. At P6 the main cell types expressing *Epas1* were ECs, astrocytes and pericytes. Among these, astrocytes were the main VEGF producers (**Extended Data Fig. 5a**) and have been reported to guide the radial expansion of the endothelial plexus^32,33^. Therefore, astrocytes could be driving the observed lesions. Reinforcing this hypothesis, it is known that astrocytes reside in the surface of the retina (**Fig. 3a**), precisely where we found HB-like lesions (**Fig. 3b,c Extended Data Fig. 5b**).

**Figure 3:**
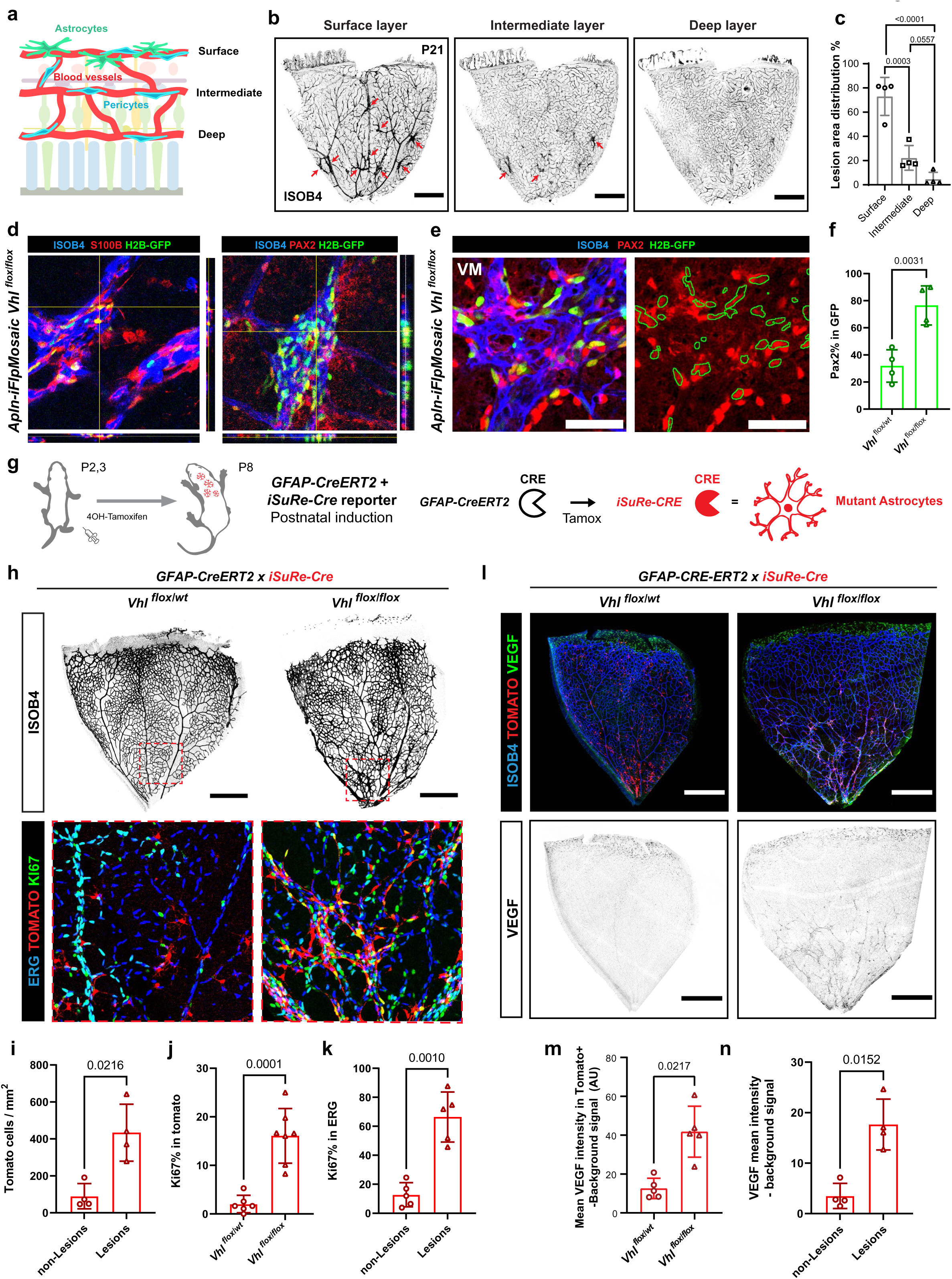
*Vhl^KO^* astrocytes induce HB-like lesions. **a,** Schematic representation of cellular distribution among the different layers of the retinal vascular plexus. Astrocytes localize in the surface layer while pericytes can be found in all layers. **b,** Representative confocal micrographs of the different vascular layers of *Apln-iFlpMosaic Vhl^flox/flox^*retinas immunostained for ISOB4. Red arrows indicate HB-like lesions. **c,** Barplot indicating the percentage of lesion distribution among the different retina vascular layers. **d,** Representative 3D confocal micrographs of *Apln-iFlpMosaic Vhl^flox/flox^* HB-like lesions immunostained for ISOB4 and the astrocyte markers S100B and PAX2 and endogenous GFP detection. **e,** Representative confocal micrograph of *Apln-iFlpMosaic Vhl^flox/flox^* HB-like lesions immunostained for ISOB4, the astrocyte marker PAX2 and endogenous GFP detection. Green outline shows the position of the GFP+ cells. **f,** Barplot indicating the percentage of astrocyte (PAX2+) cells within the GFP+ recombined cells in *Apln-iFlpMosaic* control *Vhl^flox/wt^* retinas vs *Vhl^flox/flox^* HB-like lesions. **g,** The tamoxifen-inducible and astrocyte-specific GFAP-CreERT2 and the iSuRe-Cre reporter were combined in control heterozygous (*Vhl^flox/wt^*) or homozygous (*Vhl^flox/flox^*) backgrounds to test if astrocyte-specific *Vhl* deletion could drive HB-like lesions. Mice were induced with 4-OH-Tamoxifen at P2 and P3 and dissected at P8. Recombined astrocytes are labelled with the Mb-Tomato fluorescent reporter. **h,** Representative confocal micrographs of *GFAP-CreERT2 iSuRe-Cre Vhl^flox/wt^ and Vhl^flox/flox^* retinas immunostained for ISOB4, Ki67 and endogenous Tomato detection. Magnified insets of central areas show proliferative HB-like lesions in the *GFAP-CreERT2 iSuRe-Cre Vhl^flox/flox^*. **i,** Barplot indicating the number of Tomato+ cells per mm^2^ in non-malformed vs malformed *GFAP-CreERT2 iSuRe-Cre Vhl^flox/flox^* retina areas. **j,** Barplot indicating the percentage of Ki67+ in Tomato+ recombined astrocytes in *GFAP-CreERT2 iSuRe-Cre* control *Vhl^flox/wt^* vs malformed *Vhl^flox/flox^* retinas. **k,** Barplot indicating the percentage of Ki67+ in ERG+ (endothelial) cells in non-malformed vs malformed *Apln-iFlpMosaic Vhl^flox/flox^* retina areas. **l,** Representative confocal micrographs of *Apln-iFlpMosaic Vhl^flox/wt^ and Vhl^flox/flox^*retinas immunostained for ISOB4, VEGF and endogenous Tomato detection. **m,** Barplot indicating the mean VEGF intensity within Tomato+ cells from *GFAP-CreERT2 iSuRe-Cre* control *Vhl^flox/wt^* and malformed *Vhl^flox/flox^* retinas normalized by background signal. **n,** Barplot indicating the mean VEGF intensity in non-malformed vs malformed *Apln-iFlpMosaic Vhl^flox/flox^* retina areas normalized by background signal. Data are presented as mean values +/− SD. For statistics see Source Data File 1. Scale bars are 500μm for all panels except for panel e, which is 50μm.

Immunofluorescence and 3D imaging analysis revealed that many GFP+ (*Vhl^KO^*) nuclei within the clusters were positive for the astrocytic markers S100B^34^, GFAP^35,36^ and PAX2^37^, but not for the pericyte marker DESMIN (**Fig. 3d, Extended Data Fig. 5c**). *Pax2* codes for a transcription factor labelling the nuclei of astrocytes, allowing us to confirm that 76% of the GFP+ nuclei (*Vhl^KO^*) within the lesions were positive for this astrocyte marker (**Fig. 3e,f**). This analysis suggests that *Vhl^KO^* astrocytes drive the development of retina HB-like lesions in our mouse model.

To determine if astrocyte-specific mosaic *Vhl* deletion could also lead to the development of HB-like lesions, we bred *Vhl* floxed mice with a tamoxifen-inducible astrocyte-specific CreERT2 expressing mouse model *(GFAP-CreERT2)* and the *iSuRe-Cre* allele to ensure *Vhl* deletion in all Tomato-reporter expressing cells (**Fig. 3g**). GFAP expression in retinal astrocytes has been described to start with astrocyte maturation around P2, which happens in parallel to astrocyte vascularization^33^.

Importantly, deletion of *Vhl* in GFAP+ postnatal astrocytes induced clusters and VMs (**Fig. 3h**) that closely resembled those observed in the *Apln-iFlpMosaic Vhl^flox/flox^* model (**Figs. 1 and 2**). These HB-like lesions were enriched in Tomato+ *Vhl^KO^* cells (**Fig. 3i**) and showed higher KI67% in both *Vhl^KO^* astrocytes and surrounding wildtype ECs (**Fig. 3j,k**). *Vhl^KO^* astrocytes also produced higher VEGF levels (**Fig. 3l-n**).

These results confirm that astrocyte-specific postnatal deletion of *Vhl* is sufficient to drive cluster formation, VEGF production and VMs in the murine retina, recapitulating our previous findings with the *Apln-iFlpMosaic Vhl^flox/flox^*model.

### scRNAseq analysis reveals the pathologic signature of *Vhl^KO^* astrocytes

To identify the molecular and cellular mechanisms deregulated in *Vhl^KO^*astrocytes and surrounding cells, we dissociated and antibody hashtagged retina cells from *GFAP-CreERT2 iSuRe-Cre Vhl^flox/wt^ and GFAP-CreERT2 iSuReCre Vhl^flox/flox^* P8 mice. We sorted iSuRe-Cre/Tomato+ recombined astrocytes (3% of all sorted), CD31+ ECs (25% of all sorted) and reporter negative (70% of all sorted) retina cells, that also included wildtype astrocytes (**Extended Data Fig. 6a**). scRNAseq analysis yielded 13 cell clusters with different transcriptional signatures in the two-dimensional Uniform Manifold Approximation and Projection (UMAP) (**Fig. 4a**).

**Figure 4:**
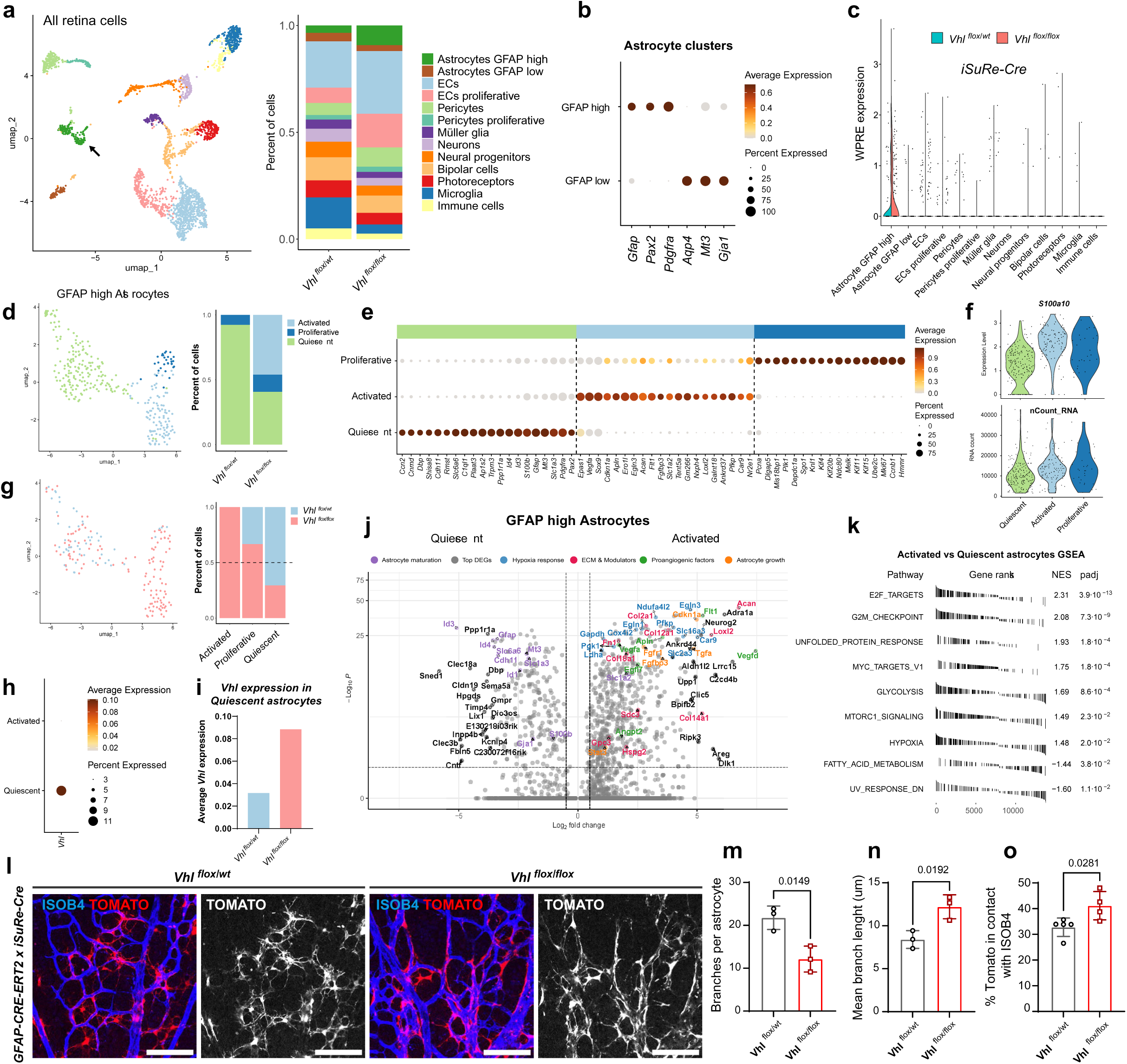
*scRNAseq* analysis of *Vhl^KO^* astrocytes reveals an activated phenotype. **a,** UMAP showing the identified retina clusters and stacked bar chart showing their relative proportions per condition. Black arrow indicates the “GFAP high” cluster. **b,** Dot plot showing the expression of key astrocyte marker genes. **c,** Violin plot showing how the expression of the iSuReCre-construct WPRE is restricted to the GFAP-high cluster. **d,** UMAP showing the identified GFAP-high astrocyte clusters and stacked bar chart showing their proportions per condition. **e,** Dot plot showing top and selected cluster marker genes for the GFAP-high astrocyte clusters. **f,** Violin plots showing the expression of the reactive A2 astrocyte marker gene, *S100a10*, and total RNA expression of the different GFAP-high astrocyte clusters. **d,** UMAP showing the distribution of *Vhl^flox/wt^* and *Vhl^flox/flox^* GFAP-high astrocyte cells (subset to the same number) and stacked bar chart showing genotype proportions per cluster. **h,** Dot plot showing average and percent expression of *Vhl* in activated and quiescent astrocyte clusters. **i,** Barplot showing the average *Vhl* expression of *Vhl^flox/wt^* and *Vhl^flox/flox^*quiescent astrocyte cells showing that *Vhl^flox/flox^* quiescent cells were not *Vhl^KO^*. **j,** Volcano plot depicting the top DEGs and other biologically significant genes when contrasting activated *Vhl ^-^* vs quiescent *Vhl ^+^* astrocyte clusters. **k,** Top significantly up and downregulated hallmark pathways in activated *Vhl ^-^*vs quiescent *Vhl ^+^* astrocyte clusters. **l,** Representative confocal micrographs of *GFAP-CreERT2 iSuRe-Cre Vhl^flox/wt^ and Vhl^flox/flox^* central areas of retinas immunostained for ISOB4 and endogenous Tomato detection showing differences in astrocyte shape. **m-o,** Barplots indicating the mean branch number per astrocyte (**m**), the mean branch lenght per astrocyte (**n**) and the percentage of Tomato+ astrocytes in contact with blood vessels (**o**) in *GFAP-CreERT2 iSuRe-Cre* control *Vhl^flox/wt^* vs malformed *Vhl^flox/flox^* retinas. Data are presented as mean values +/− SD. For statistics see Source Data File 1. Scale bars are 100μm.

Clusters were annotated according to the differential expression of already established cell-type specific markers^22^ (**Extended Data Fig. 6b**). We identified two populations of retina astrocytes, as previously described^38,39^. One was characterized by high *Gfap*, *Pax2* and *Pdgfra* expression and the other by low *Gfap* expression and high expression of astrocyte endfeet markers like *Aqp4, Mt3* and *Gja1* (**Fig. 4b** and **Extended Data Fig. 6b,c**). Analysis of the recombined *iSuRe-Cre* reporter expression (WPRE sequence) confirmed its expression mostly by the *Gfap* high astrocytes (**Fig. 4c**). Reclustering of the *Gfap* high astrocytes yielded 3 cell clusters (**Fig.4d)**. These were defined by the expression of specific marker genes that indicated a quiescent, activated or proliferative state (**Fig. 4e and Extended Data Fig. 7a,b**). In line with this, the cells within the activated and proliferative clusters expressed the reactive astrocyte (A2) marker S100a10^40^ and showed higher transcriptomic activity (**Fig.4f)**. Interestingly, the “activated” cell cluster was exclusively present in the *Vhl^flox/flox^* sample (**Fig. 4g**). This cluster had complete loss of *Vhl* expression (**Fig. 4h**). Notably, the quiescent astrocytes found within the *Vhl^flox/flox^* sample had even higher expression of *Vhl* than in the *Vhl^flox/wt^* sample (**Fig. 4i**), suggesting these were mostly *iSuRe-Cre* negative, wildtype astrocytes, that as mentioned before were also co-sorted.

Given that the “activated” cell cluster represented *Vhl^KO^*GFAP-high astrocytes, we next analysed in depth the genes differentially expressed (DEGs) by this cluster (**Fig. 4j**). *Vhl^KO^* Astrocytes showed upregulation of hypoxia-pathway related genes such as the prolyl hydroxylases genes *Egln3* and *Egln1*^41^, respiratory complex IV paralog subunits like *Ndufa4l2*^42,43^ *and Cox4i2*^44^, and *Car9*. *Car9* encodes for the Carbonic Anhydrase IX, an enzyme that is highly upregulated upon VHL deletion and commonly used in the histopathological diagnosis of VHL disease^45^. The upregulation of these genes is likely a direct effect of *Vhl* loss and HIFs stabilization. In addition, genes related with hypoxia-induced metabolic activation of glycolytic metabolism like *Ldha*, *Slc16a3*, *Pfkp*, *Pgk1*, *Slc2a3* and *Gapdh* ^46–48^ were strongly upregulated too. *Vhl^KO^* astrocytes also expressed higher levels of angiogenic molecules, such as *Vegfa, Vegfd, Angpt2, Tgfb1*^49^ and *Apln*^50^ and also extracellular matrix remodelling genes, such as *Fn1, Loxl2*, *Col12a1* previously linked to retina neovascularization^51,52^. Interestingly, *Acan*, a Chondroitin Sulfate Proteoglycan (CSPG) commonly produced by activated A2 astrocytes^40,53^ was a top upregulated gene in the “activated” astrocyte population. Astrocyte growth factor related genes like *Fgfr1, Gfgbp3, Stat3*^54,55^ or *Tgfa*, a strong promoter of astrocyte proliferation *in vivo* ^56,57^, were also among the highest DEGs. Besides the higher expression of markers related with increased cellular and metabolic activity, *Vhl^KO^* astrocytes also showed a strong downregulation of markers related with mature astrocytes, such as the transcription factors *Id3*, *Id4* and *Id1*^58^; and astrocyte structural proteins like *Gfap, S100b, Mt3, Slc1a3 and Gja*^39,55^ (**Fig. 4j**). GFAP downregulation was confirmed by retina immunostaining (**Extended Data Fig. 7c**). Hallmark and Gene Set Enrichment Analysis (GSEA) confirmed an enrichment in cell cycle and cell growth pathways (E2F_TARGETS, G2M_CHECKPOINT, MYC_TARGETS_ and MTORC1_SIGNALING) and increased expression of genes related with hypoxia, glycolysis and the unfolded protein response (**Fig. 4k**). Conversely the FATTY_ACID_OXIDATION hallmark was downregulated, confirming the shift in metabolism towards glycolysis.

Besides the analysis of DEGs, the imaging of the *iSuRe-Cre* membrane Tomato reporter showed that *Vhl^KO^* astrocytes lost the typical star shape of astrocytes, and instead they were in general less ramified and more fusiform (**Fig. 4l-n**). This fusiform shape has been previously described in gliosis and other pathophysiological processes involving activated astrocytes^59,60^. *Vhl^KO^* astrocytes were also closer and more frequently wrapping around blood vessels, rather than contacting them through their end feet. As a result, astrocyte-endothelial contact area was increased (**Fig. 4l,o**). This higher cell surface contact may increase their molecular crosstalk with blood vessels, which can be a response to the genetic hypoxia and reprogramming experienced by *Vhl^KO^* cells.

Overall, these results suggest that *Vhl* deletion in GFAP+ astrocytes increases the hypoxia and cellular growth signalling leading to an abnormal activated cell state, impairing their proper maturation, and inducing the abnormalization of the surrounding endothelium.

### Astrocyte *Vhl* deletion enhances their crosstalk with ECs inhibiting their maturation

To understand how *Vhl* deletion in astrocytes changes their molecular communication with ECs, we used the scRNAseq data to evaluate changes in the receptor-ligand expression and interactions between astrocytes and ECs (**Extended Data Fig. 8a**) through CellChat analysis^61,62^. Interestingly, both the total number and strength of receptor-ligand interactions were 4 times higher in the *Vhl^flox/flox^*retinas (**Fig. 5a**). In the *Vhl^flox/flox^* retinas, “activated” and “proliferative” astrocytes were the clusters with the highest counts of established outgoing interactions (**Fig. 5b**). We found a significant enrichment of proangiogenic signalling pathways including ANGPT, ANGPTL, COLLAGEN, VEGF, FN1, MK, APELIN and PTN. In addition we also observed enrichment of pathways related with astrocyte migration and patterning like LAMININ^63^, PDGF^52,64,65^ or NCAM, some of which have also been reported to play a role in physiological and pathological angiogenesis^66^ (**Fig. 5c and Extended Data Fig. 8b,c**). Regarding the directionality and strength of these interactions, the signals from “activated” and “proliferative” astrocytes to ECs were the strongest ones, followed by astrocyte-to-astrocyte interactions, which were also enhanced (**Fig. 5d,e**). Ligand-receptor pair analysis showed that the “proliferative” and “activated” *Vhl^KO^* astrocytes had an increased communication with ECs trough Vegfa-to-Vegfr2/Vegfr1, Fn1-to-Itga4/5/Itgb1 and Col2a1-to-Itga1/Itgb1 (**Fig. 5f**).

**Figure 5:**
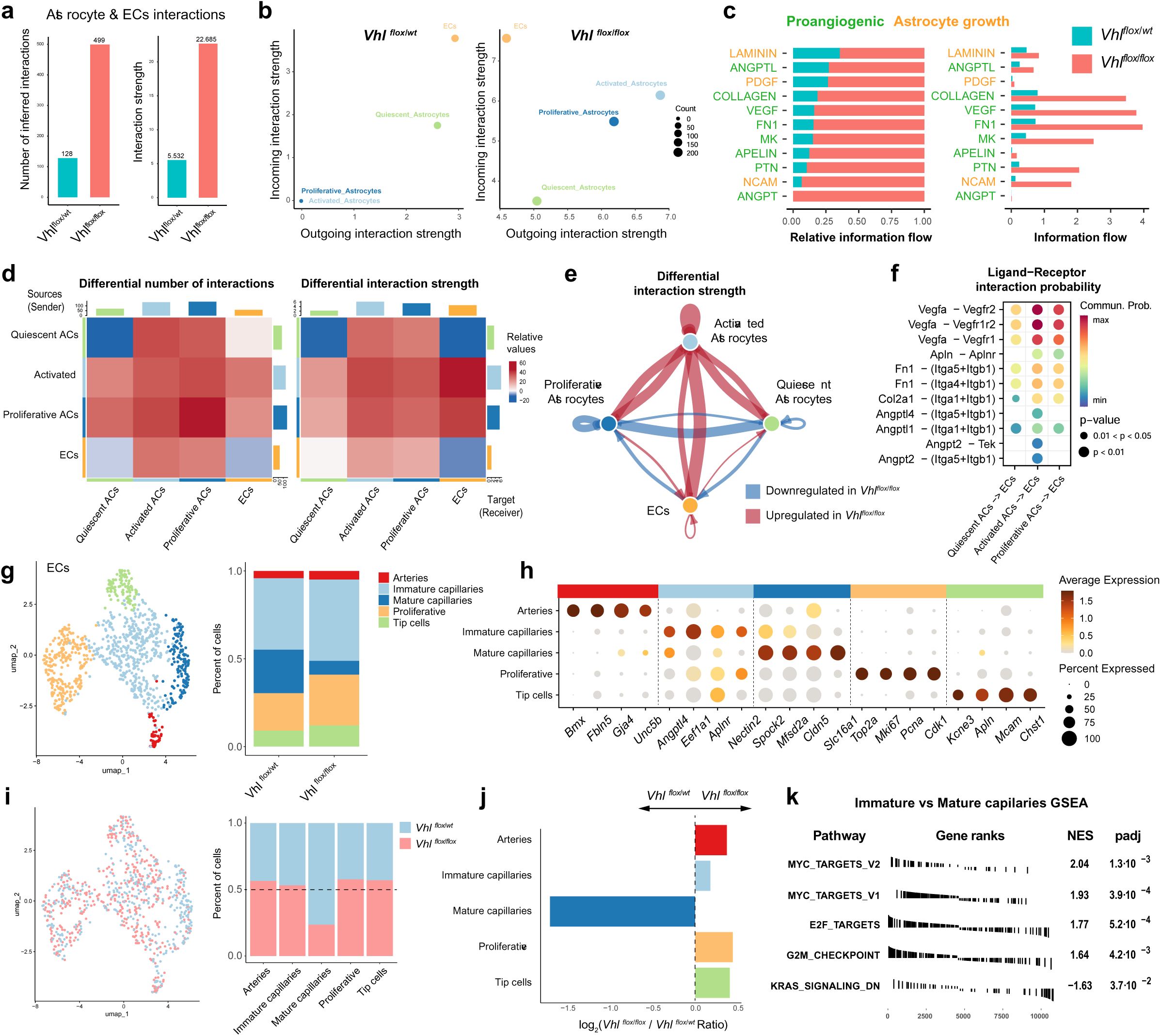
Astrocyte-Endothelial signalling and endothelial characterization. **a,** Barplots showing the total number of interactions and interaction strength of the inferred cell-cell communication networks from *Vhl^flox/wt^* vs *Vhl^flox/flox^* GFAP-high astrocytes and ECs. **b,** Scatterplot comparing the outgoing and incoming interaction strength of the indicated clusters and genotypes in 2D space. **c,** Stacked and histogram bar graphs representing relative and absolute information flow of the signalling pathways enriched in the *Vhl^flox/flox^*condition. Green pathways are related with proangiogenic pathways, orange with astrocyte growth. **d,** Heatmap showing differential number of interactions and interaction strength. The top colored bar plot represents the sum of incoming signaling relative values. The right colored bar plot represents the sum of outgoing signaling relative values. Red or blue represents increased or decreased signalling, respectively, in the *Vhl^flox/flox^*dataset compared to the *Vhl^flox/wt^* one. **e,** Circle plot representing the differential interaction strength between *Vhl^flox/flox^*and *Vhl^flox/wt^* conditions (red is upregulated in *Vhl^flox/flox^,* blue is downregulated in *Vhl^flox/flox^*). **f,** Bubble plot comparing the highest communication probabilities mediated by ligand-receptor pairs from the different astrocyte clusters to ECs in the *Vhl^flox/flox^* sample **g,** UMAP showing the identified endothelial clusters and stacked bar chart showing their proportions per condition. **h.** Dot plot showing top and selected cluster marker genes for the endothelial clusters. **i,** UMAP showing the distribution of *Vhl^flox/wt^*and *Vhl^flox/flox^* endothelial cells (subset to the same number) and stacked bar chart showing genotype proportions per cluster. **j,** Histogram bar plot represents the log2 ratios of *Vhl^flox/flox^* over *Vhl^flox/wt^* cells for each endothelial cluster. **k,** Top significantly up and downregulated hallmark pathways in Immature vs Mature capillary clusters.

These results suggest that the “activated” *Vhl^KO^* astrocytes are proangiogenic to neighbouring ECs, stimulating their growth in a paracrine fashion. This is consistent with our previous analysis showing higher VEGF expression in mutant astrocytes (**Fig. 2**), and the decrease in lesions after VEGF trap or Epas1 loss-of-function (**Extended Data Fig.3,4**).

To better understand the changes happening in the endothelial population we performed subsetting and higher resolution analysis of the endothelial scRNAseq data obtained from *Vhl^flox/wt^* and *Vhl^flox/flox^* retinas. This revealed the existence of 5 clearly defined clusters that we annotated as arteries, immature capillaries, mature capillaries, proliferative and tip cells (**Fig. 5g**). Annotation and validation of these clusters was performed using marker genes previously reported^22^(**Fig. 5h**). Analysis of the different populations showed an increase in the proportion of proliferative, immature and tip ECs and a decrease in the frequency of mature or quiescent ECs in *Vhl^flox/flox^* retinas (**Fig. 5i,j**), which is in line with our previous histological analysis (**Fig. 3h**). Note that only a relatively small subset of vessels within the *GFAP-CreERT2, iSuRe-Cre, Vhl^flox/flox^* retinas became malformed and angiogenic (**Fig. 3h**), and this scRNAseq data of all collected ECs is in line with this observation. GSEA analysis showed that the immature capillaries were enriched in cell growth or activation pathways like MYC_TARGETS_V1 and V2, E2F_TARGETS and G2M-CHECKPOINT (**Fig.5-k).** These pathways’ upregulation could prevent capillary maturation, leading to VMs development.

Overall, these results indicate that *Vhl^KO^* “activated” astrocytes increase their molecular interactions with ECs and are proangiogenic, preventing endothelial maturation and leading to VMs development.

### *Vhl* deletion in adult mice does not induce hemangioblastomas

In VHL patients, new HBs arise during the adolescence and adult life. Pharmacological management of VHL disease HBs aims to prevent the appearance or growth of lesions. However, it is still not clear if HBs are originated by de-novo somatic mutations in adults, or if they arise from latent precursor lesions that originated during organ development and become reactivated after^13^, as it happens in other types of VMs^67^.

To answer this question, we induced the mosaic deletion of *Vhl* in postnatal and adult mice having the ubiquitously expressed *CAG-FlpOERT2* and *iFlpMosaic* alleles (**Extended Data Fig. 9a**) or the astrocyte-specific *GFAP-CreERT2* and *iSuRe-Cre* alleles (**Extended Data Fig. 9b**). Both the ubiquitous and astrocyte-specific inducible models presented hemangioblastomas when induced postnatally (**Extended Data Fig. 9c-f, left panels),** but not when induced in adult mice (**Extended Data Fig. 9c,d right panels)**. Interestingly, even when ubiquitously induced, postnatal *CAG-FlpOERT2 iFlpMosaic* lesions were also enriched in *Vhl^KO^* PAX2+ astrocytes (**Extended Data Fig. 9g,h**), highlighting again the role of astrocytes in driving hemangioblastoma lesions after *Vhl* deletion.

We then wondered if the lesions induced postnatally would continue to proliferate for longer periods of time into early adulthood. For that we assessed VMs activation by KI67 detection. Overall, most lesions became quiescent from P7 to P21 (**Extended Data Fig. 10a,b**), with very few showing KI67 signals (**Extended Data Fig. 10a right panels)**. This data suggests that there is a specific time-window for single mutant astrocytes to grow and develop into clusters able to induce VMs. After angiogenesis is finished and the tissue reaches its maturation, the large majority of *Vhl^KO^* cells and surrounding vessels no longer proliferate. However, it is possible they become reactivated after tissue injury or additional inflammatory signals.

### Rapamycin treatment prevents hemangioblastoma development after loss of *Vhl*

In our scRNAseq analysis, mTOR signalling showed among the top upregulated pathways in the activated *Vhl^KO^* astrocytes (**Fig. 4m**). Given the extended use of the FDA-approved drug rapamycin (mTOR inhibitor) in the clinics and as a treatment for other VMs^68,69^, we wondered if this drug could also be used to prevent the development of *Vhl^KO^* clusters and associated VMs in our model. For this experiment, GFAP-CreERT2 *Vhl^flox/wt^* and *Vhl^flox/flox^* mice were induced with 4OHT at P2 and P3. Rapamycin (4mg/kg) or vehicle were administrated at P5, P6 and P7. Mice were dissected at P8 (**Fig. 6a**). Rapamycin significantly reduced KI67 frequency in ECs, EC density and astrocyte proliferation (**Fig. 6b-e**), with no VMs or hemangioblastomas. Rapamycin also reduced expression of P21, but did not change the expression of VEGF per *Vhl^KO^*Tomato+ astrocyte (**Fig. 6f-h**). This suggests that even though rapamycin can reduce the activated state and proliferation of *Vhl^KO^* Tomato+ astrocytes, it cannot prevent their genetic hypoxia and VEGF expression. However, since it reduces their clonal expansion and clusterization, it can effectively reduce the total amount of astrocytes producing VEGF in a single vascular area (**Fig. 6i**). In addition, we found *Vhl^KO^* astrocyte morphology to be more normal and similar to controls after rapamycin treatment (**Fig. 6j-l**).

**Figure 6:**
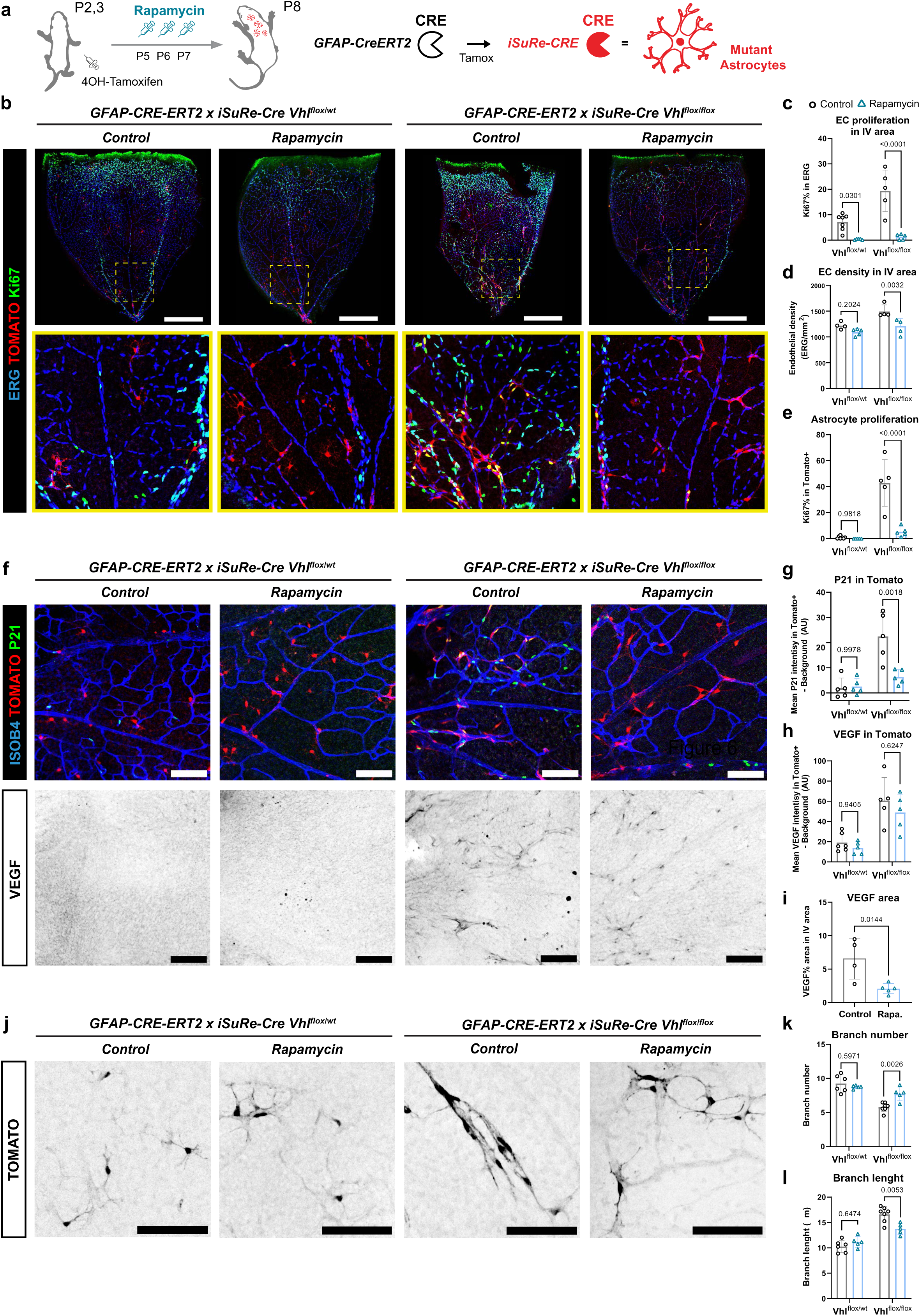
Rapamycin treatment reverts *Vhl^KO^* astrocyte activation and HB-like lesion development. **a,** Treatment schedule: *GFAP-CreERT2 iSuRe-Cre Vhl^flox/wt^* and *Vhl^flox/flox^* mice were induced with 4-OH-Tamoxifen at P2 and P3 and rapamycin (4mg/kg) was injected IP at P5,6 and 7. Mice were dissected at P8. **b**, Representative confocal micrographs of control and rapamycin treated *GFAP-CreERT2 iSuRe-Cre Vhl^flox/wt^ and Vhl^flox/flox^*retinas immunostained for ERG, Ki67 and endogenous Tomato detection. Magnified insets of central intervein (IV) areas show a reduction in proliferative lesions after rapamycin treatment. **c-e,** Barplots showing rapamycin (blue) effect on endothelial proliferation (Ki67% in ERG+ cells) in the intervein central areas of the retina (**c**), on endothelial density (ERG+ cells / mm^2^) in the intervein central areas of the retina (**d**), and on astrocyte proliferation (Ki67+ cells / Tomato+ cells, **e**)**. f,** Representative confocal micrographs of control and rapamycin treated *GFAP-CreERT2 iSuRe-Cre Vhl^flox/wt^ and Vhl^flox/flox^* retinas immunostained for ISOB4, P21, VEGF and endogenous Tomato detection. **g,** Barplots showing rapamycin effect (blue) on the activated astrocyte marker P21 expression (P21+ cells / Tomato+ cells). **h,** Barplots showing that rapamycin has no effect (blue) on single cell VEGF production (Mean VEGF expression in Tomato+ / Background signal). **i,** Barplots showing rapamycin effect (blue) on high VEGF area (VEGF area / Intervein area). **j,** Representative confocal micrographs of control and rapamycin treated *GFAP-CreERT2 iSuRe-Cre Vhl^flox/wt^ and Vhl^flox/flox^* retinas, endogenous Tomato detection. **k,** Barplots showing rapamycin effect (blue) on astrocyte branch number. **l,** Barplots showing rapamycin effect (blue) on astrocyte branch length. Data are presented as mean values +/− SD. For statistics see Source Data File 1. Scale bars are: b-500 μm; f,j-100μm. Abbreviations: P2,3,5,6,7 – Postnatal day 2,3,5,6,7; Tamox – Tamoxifen; IV – Intervein.

Collectively, these findings suggest that rapamycin treatment reduces *Vhl^KO^* astrocyte activation and proliferation, without affecting their genetic hypoxia. The reduction in their expansion and clusterization, leads to less *Vhl^KO^* cells producing VEGF, and consequently, less VMs development. The additional effect of rapamycin in blocking endothelial proliferation, further reduces the development of hemangioblastomas. This can be a very interesting alternative to treat VHL disease hemangioblastomas, as the use of the HIF-2A inhibitor belzutifan has significant side-effects^4^ and should only target the *Vhl^KO^* cells and not the existing VMs.

## Discussion

Hemangioblastoma treatment mostly relies on invasive surgical interventions that can negatively impact patients’ life expectancy, especially in the context of VHL disease, where hemangioblastoma recurrence implies frequent neuro-surgical interventions^3^. The scarce pharmacological options for VHL disease treatment ^4^ result from the limited understanding of key aspects of hemangioblastoma development, such as the cellular origin of the disease, the mechanisms driving the abnormal vascularization, the timing of VHL mutation and disease onset, or the mechanisms behind VHL-lesions reactivation and recurrence.

Animal models with induced loss of VHL function were used in the past^11,12,70–72^. However, none allowed the induction of traceable genetic mosaics of *Vhl^KO^* and *Vhl^WT^* cells. Which is of particular interest considering the mosaic origin of VHL disease. Here, we used iFlpMosaics^14^ to induce and quantify with very high precision how *Vhl^KO^* mutant cells behave during the postnatal and adult stages.

Following the hemangioblast origin hypothesis proposed previously^8,73–75^, and the expression of *Apln* in hemangioblasts, we induced *iFlpMosaics* with *Apln-FlpO*. This resulted in hemangioblastoma-like lesions in the central nervous system, containing both proliferative *Vhl^KO^* non-ECs surrounded by *Vhl^WT^*malformed vessels, closely resembling human hemangioblastomas where the tumour compartment is *VHL^KO^* but the vasculature remains *VHL^WT^*^5,6,26,27^. We showed that these clusters of *VHL^KO^* mutant cells did arise from proliferation of *Apln*-derived non-ECs, that we found to be astrocytes. We also saw that hemangioblastomas are formed when vessels start to mature and VEGF levels are relatively low. Our work indicates that *Vhl^KO^*cells become malignant by inhibiting the normal vascular maturation process, that requires a switch from an angiogenic to a quiescent state, a process that is impaired when there are clusters of many VEGF-producing *Vhl^KO^*cells in the vicinity of blood vessels undergoing maturation.

Mechanistically, the deletion of *Epas1* (HIF-2α) was sufficient to completely prevent VEGF expression by astrocytes, and the appearance of VMs. *Epas1* deletion also reduced the expansion and clusterization of *Vhl^KO^* astrocytes, confirming it to be the key HIF modulated by VHL in astrocytes. Therefore, this work provides mechanistic insight into the reported efficacy of Belzutifan, a HIF-2α inhibitor that is so far the only approved pharmacological treatment against VHL disease hemangioblastomas ^4^. It is known that VHL regulates both HIF-2α and HIF-1α, being the latter the one most frequently associated to the regulation of VEGF expression^29^. However, we found that in astrocytes, only HIF-2α, not HIF-1α, was relevant for the biology and pathogenesis of *VHL^KO^* cells. This differential effect of HIF-1α and HIF-2α is in line with previously published data showing that HIF-1α has a tumour suppressor role in clear cell renal cell carcinoma, which contrasts with the oncogenic role of HIF-2α^76–78^.

Our work indicates that astrocytes can drive VHL-hemangioblastomas in the central nervous system. By using the *GFAP-CreERT2* and *iSuRe-Cre* alleles, we could profile by scRNAseq *Vhl^KO^*astrocytes for the first time, and in this way identify the molecular mechanisms that lead to their pathogenesis. This analysis revealed that *Vhl^KO^*astrocytes upregulate many genes related with genetic hypoxia, metabolic rewirement (decrease in Fatty acid oxidation and increase in glycolysis and lactic fermentation), proliferation and cell growth (E2F, G2M_Checkpoint, M-TORC1 and MYC pathways), which explain the rapid formation of large astrocytes clusters from few mutant cells between P3 and P7. In addition, *Vhl^KO^* astrocytes expressed several ECM molecules and growth factors like VEGF, that overall induce angiogenesis and impair vascular maturation. CellChat analysis^62^ identified activated astrocytes as signalling hubs towards ECs, with several enriched pathways, like Angiopoietin, PTN or VEGF. The analysis of single *Vhl^KO^*astrocytes shape also revealed that they lose the typical star shape of mature astrocytes^59^, and instead adopt a more fusiform shape typical of immature astrocytes, that further increases the surface of interaction with ECs, boosting their crosstalk and molecular influence on ECs. Interestingly, both the morphological structure and molecular profiles of *Vhl^KO^* astrocytes are very similar to the ones reported on reactive A2 astrocytes found after ischemic injury^40,53^.

An important finding of our study is the limited temporal window for hemangioblastomas development after the loss of *Vhl* function. We found that, in the retina, lesions could only initiate between P3 and P21, and only in the vascular areas undergoing maturation. The induction of the mutation in adult stages did not cause hemangioblastomas in the central nervous system. This suggests that there is a temporal window in which both the *Vhl^KO^* astrocytes and surrounding ECs are still plastic and strongly influenced by the mutation. Later, after the tissue becomes quiescent, cells no longer proliferate and form *de novo* lesions. This data supports the already suggested idea that VHL disease patients harbour early subclinical precursor lesions^13,79^, as the ones identified in this study, that later in life can become more severe. The latency of the lesions could be broken by hormonal changes, inflammation, or surgical trauma. Indeed, other diseases involving vascular lesions like arteriovenous malformations can also be latent until reactivated by sporadic events^80^.

Given that rapamycin was able to effectively reduce the growth and clusterization of *Vhl^KO^* astrocytes and ECs, effectively preventing the development of lesions, we propose the use of Rapamycin (which targets both the *Vhl^KO^* cells and ECs) alone or in combination with the HIF-2α inhibitor Belzutifan^4^ to enhance its therapeutic efficacy. Since Belzutifan treatment adherence is limited by strong side-effects like anaemia, treatment with Rapamycin (as used in some VMs^68^) could be less toxic, or allow dose reduction of Belzutifan, while maintaining efficacy, potentially improving both life expectancy and quality of life for VHL patients.

Given the many deregulated genes and molecular signatures identified in our *Vhl^KO^*scRNAseq analysis, it may be now possible to identify even more specific molecular targets and drugs in the future.

In conclusion, our findings identify *Vhl^KO^* astrocytes as central drivers of hemangioblastomas associated with VHL disease and provide new mouse models and molecular insights to support future studies aimed at understanding and targeting this disease.

## Acknowledgments

Research in Rui Benedito’s laboratory was supported by the European Research Council (ERC) Consolidator Grant AngioUnrestUHD (101001814), the Ministerio de Ciencia, Innovación y Universidades (PID2020-120252RB-I00 and PID2023-148880OB-I00), and “la Caixa” Banking Foundation (HR22-00316 AngioHeart), all awarded to Rui Benedito. Macarena de Andrés-Laguillo was funded by the Ministerio de Ciencia, Innovación y Universidades (PRE2019-087459) and by the Instituto de Salud Carlos III, through the Complementary Actions for International Joint Programming 2023, within the project entitled “Bridging the gap between cardiac and vascular regeneration” (acronym: RESCUE), ref.: AC23_2/00019. This project has received funding from the Instituto de Salud Carlos III (ISCIII) and has been co-funded by the European Union under the Horizon Europe Framework Programme, grant agreement No. 10105426.

J.A.E. is supported by RTI2018-099357-B-I00, PID2021-1279880B, and TED2021-131611B-I00 funded by MCIN/AEI/10.13039/501100011033 and the European Union “NextGenerationEU”/Plan de Recuperación Transformación y Resiliencia/PRTR; CIBERFES (CB16/10/00282); Fundación “la Caixa” (LCF/PR/HR23/52430010). K.D.B. is endowed by the Schulthess foundation. A.D and L.C were partially supported by Comunidad de Madrid (S2022/BMD-7403 RENIM-CM) and by Instituto de Salud Carlos III (PT20/00044), cofunded by European Union, European Regional Development Fund (ERDF, “A way of making Europe”). ReDIB ICTS infrastructure TRIMA@CNIC, Ministerio de Ciencia e Innovación (MCIN).

The CNIC is supported by the Instituto de Salud Carlos III (ISCIII), the Ministerio de Ciencia, Innovación y Universidades (MICIU), and the Pro CNIC Foundation. It is recognized as a Severo Ochoa Center of Excellence (grant CEX2020-001041-S funded by MICIU/AEI/10.13039/501100011033).

Microscopy experiments were carried out at the CNIC Microscopy and Dynamic Imaging Unit, an ICTS-ReDib facility co-funded by MCIN (/AEI/10.13039/501100011033) and the ERDF “A way to build Europe” (#ICTS-2018-04-CNIC-16).

We thank the members of the CNIC microscopy, genomics, advanced imaging, cytometry, and bioinformatics units for their support throughout the project. We thank the Dr. Silvia Martín-Puig for sharing the *Vhl^flox^, Epas1^flox^ and Hif1a^flox^* models. We thank the Prof. Juan Pedro Bolanos and the Dr. Eduardo Balsa for sharing the GFAP-CreERT2 model. We thank the Neurovascular Pathophysiology Lab led by Professor Mª Ángeles Moro for sharing reagents and for their valuable advice.

## Authors contribution statement

M.dA-L and R.B. designed the experiments, interpreted the results, assembled the figures, and wrote the manuscript. M.dA-L performed most experimental procedures, conducted the scRNA-seq bioinformatic analysis, wrote and edited the text, and assembled the figures. I.G-G. developed the iFlpMosaic genetic tools used in the study. S.F-R gave support with manuscript edition. A.G-C. genotyped the mouse colonies. S.R-G helped with colony management and mice dissection. L.D-D helped with whole organ tissue clearing and image acquisition. A.D performed the design, manufacturing, and chemical validation of the nanoparticles used for the MCT. L.C selected the contrast agent, defined the imaging protocol and performed image post-processing. JA.E. and K.D.B. provided scientific guidance and contributed to data interpretation. All authors approved the final version of the manuscript.

## Extended Data Figure Legends

**Extended Data Figure 1:**
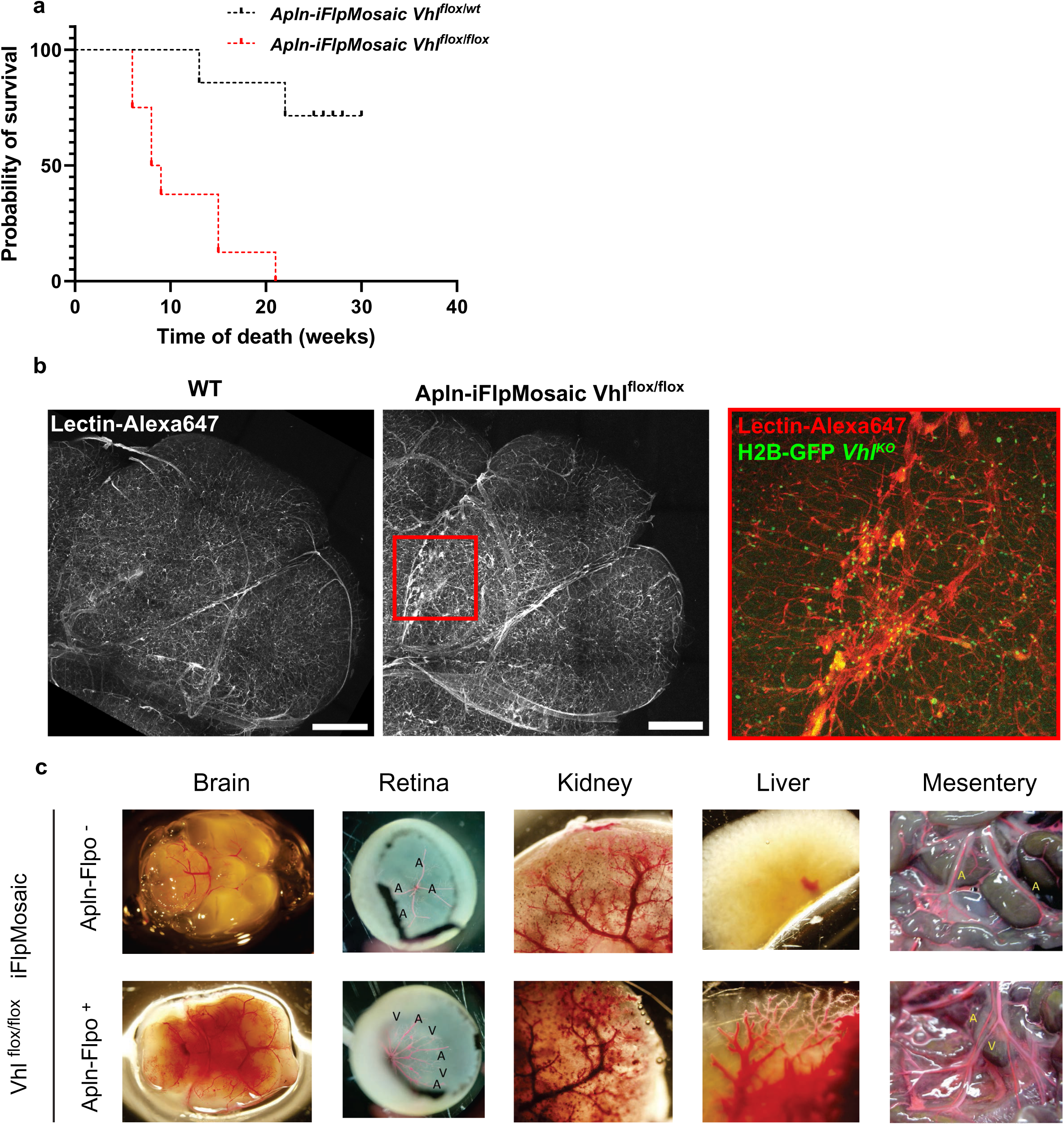
Characterization of *Apln-iFlpMosaic Vhl^flox/flox^* adult mice. **a,** Survival curve of *Apln-iFlpMosaic* control *Vhl^flox/wt^* vs *Vhl^flox/flox^*. Median survival of *Vhl^flox/flox^* is 8.5 weeks. **b,** Representative confocal 3D stacked micrographs of wild type and *Apln-iFlpMosaic Vhl^flox/flox^*cerebellums perfused with fluorescent labelled lectin (Lectin-Alexa647). Endogenous GFP detection labels *Vhl^KO^*cells. **c,** Representative pictures from left ventricle latex perfused control and *Apln-iFlpMosaic Vhl^flox/flox^*organs. Brain, kidney and liver were cleared with CUBIC1. For statistics see Source Data File 1. Scale bars are 500μm.

**Extended Data Figure 2:**
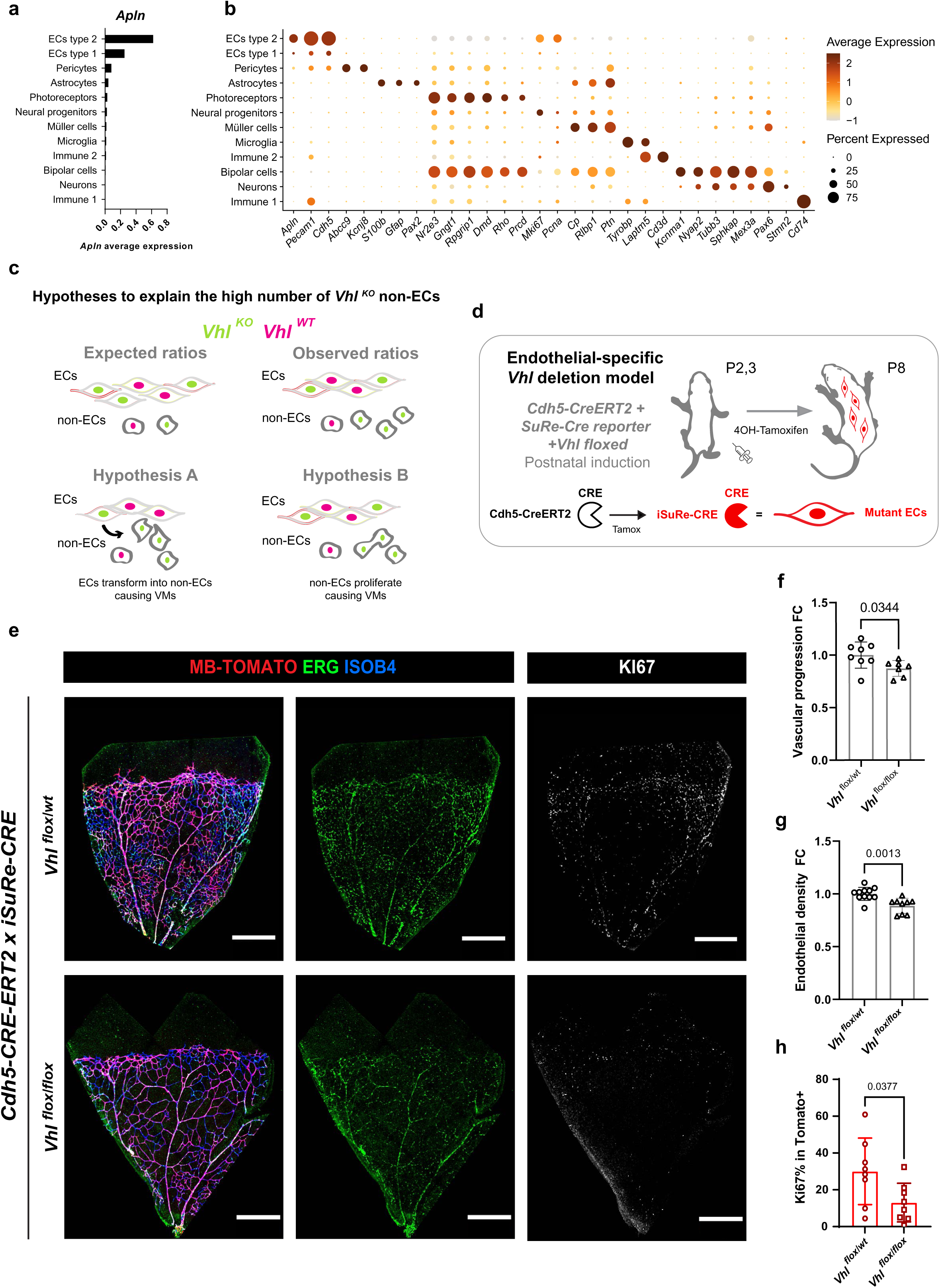
Apln-derived *Vhl^KO^* ECs don’t drive HB-like lesions. **a,** Barplot showing mean *Apln* expression in the different postnatal retina cell types. **b,** Dot plot showing the expression of the different cell type markers. **c,** Graphical scheme showing how, in the *Apln-iFlpMosaic Vhl^flox/flox^*retinas, tzahe number of *Vhl^KO^* non-ECs is higher than the one that would be expected in a control mosaic. Different hypotheses could explain this fact: A, *Vhl^KO^* ECs transdifferentiate into non-ECs; B, some *Vhl^KO^* non-ECs expand via proliferation. **d,** *Cdh5*-*CreERT2* with the *iSuRe-Cre* reporter and the floxed *Vhl* allele, which ensures Cre-recombination in any tomato+ EC were induced at P2 and P3, retinas were dissected at P8. **e,** Representative confocal micrographs of *Cdh5*-*CreERT2* iSuRe-Cre *Vhl^flox/wt^* and *Vhl^flox/flox^* retinas immunostained for ISOB4, ERG, KI67 and endogenous detection of Tomato showing decreased angiogenesis but no HB-like lesions development. **f,** Barplot showing vascular progression normalizing (fold change) each value by the average vascular progression from the littermate controls. **g**, Barplot showing vascular density (ISOB4 area in total vascularized retina area) normalizing (fold change) each value by the average vascular density from the littermate controls **h,** Barplot showing the percentage of KI67+ cells within *Vhl^flox/wt^* and *Vhl^flox/flox^* recombined ECs. Data are presented as mean values +/− SD. For statistics see Source Data File 1. Scale bars are 500μm. Abbreviations: P – Postnatal day. Single cell RNAseq data was reanalysed from Zarkada et.al.

**Extended Data Figure 3:**
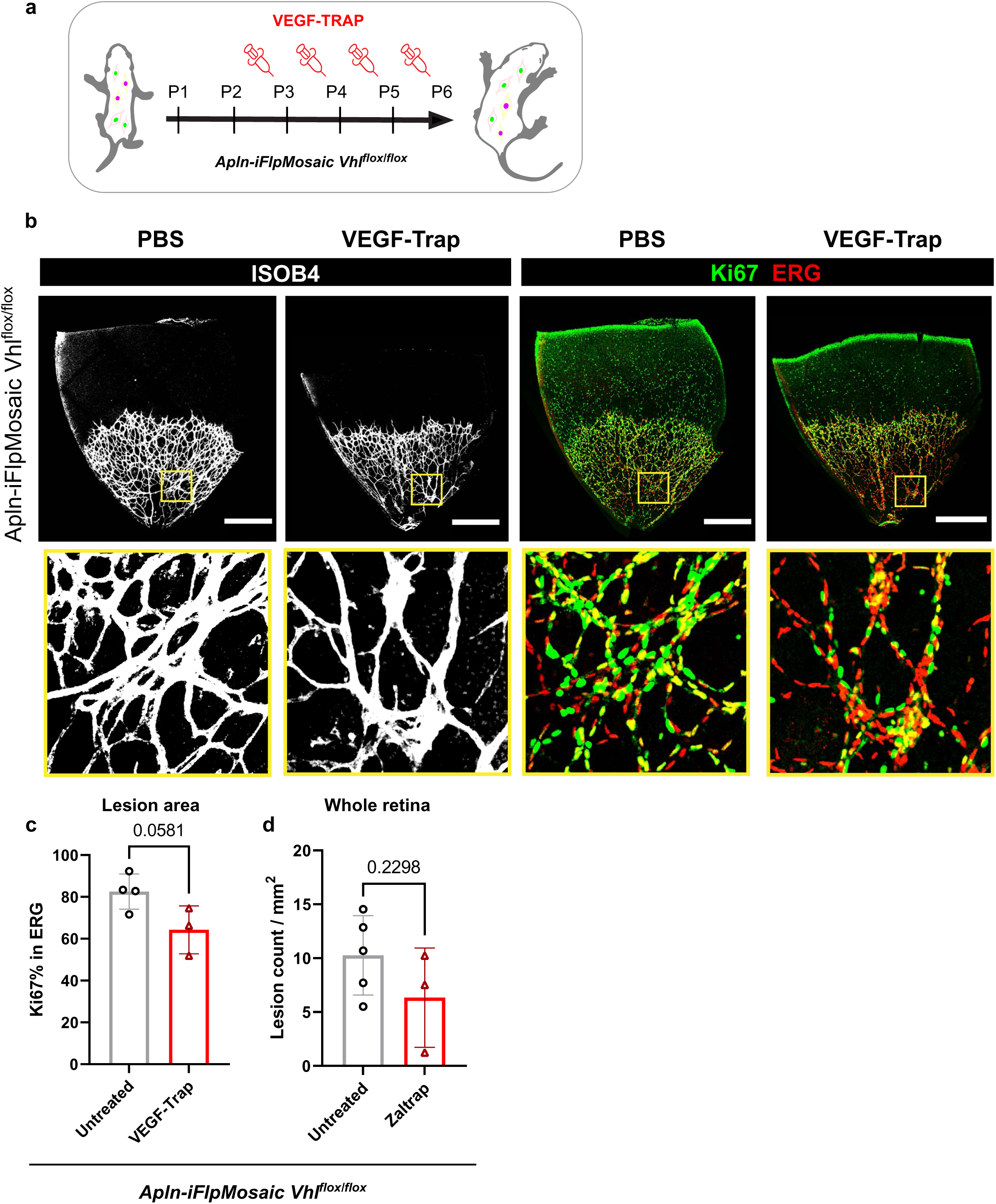
VEGF trap treatment with Zaltrap. **a,** Apln-iFlpMosaic *Vhl^flox/flox^* mice were treated with 20mg/kg Zaltrap at P3,4,5,6 and dissected at P6. **b,** Representative confocal micrographs of PBS and Zaltrap treated *Apln-iFlpMosaic Vhl^flox/flox^*retinas immunostained for ISOB4, ERG and KI67. **c,** Barplot showing the percentage of KI67+ ECs (ERG+ cells) in the untreated vs Zaltrat treated *Apln-iFlpMosaic Vhl^flox(flox^* retinas. **b,** Barplot showing the VMs count per mm^2^ in the untreated vs Zaltrat treated *Apln-iFlpMosaic Vhl^flox(flox^* retinas. Data are presented as mean values +/− SD. For statistics see Source Data File 1. Scale bars are 500μm. Abbreviations: P – Postnatal day.

**Extended Data Figure 4:**
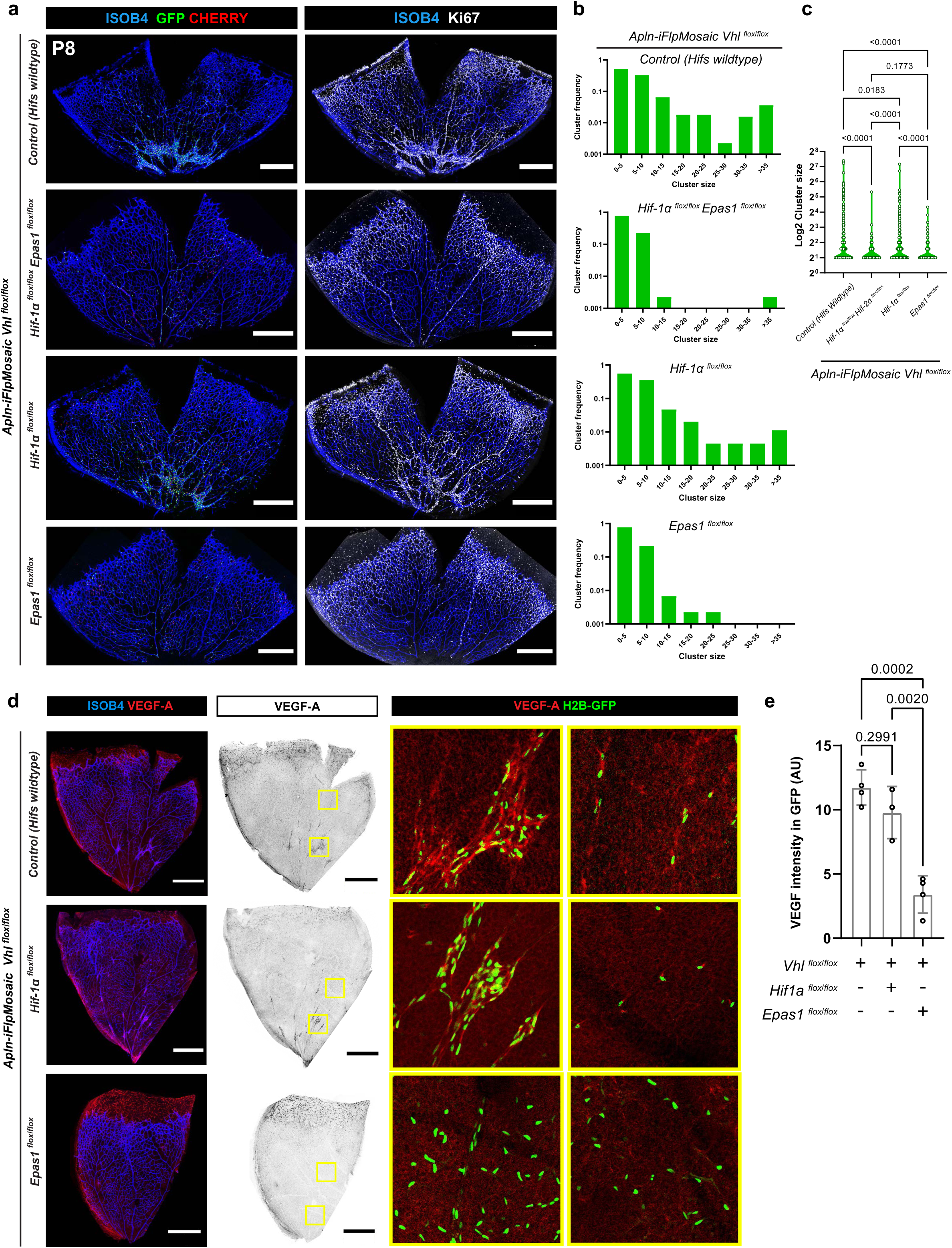
*Epas1,* but not *Hif1a* codeletion fully rescues the *Apln-iFlpMosaic Vhl^flox/flox^* phenotype. **a,** Representative confocal micrographs of *Apln-iFlpMosaic Vhl^flox/flox^* retinas with codeletion of the indicated HIF-encoding-genes, showing how HB-like lesions depend on Epas1 expression to be formed. Retinas were immunostained for ISOB4 and KI67, endogenous Cherry and GFP detection. **b,** Histograms showing the GFP (mutant) cluster frequency (y axis) and size (x axis) from a total of 445 clusters from 4 independent mice for each genotype. Samse number of clusters were analysed for each condition. Individual cells were not considered as clusters **c,** Violin plot showing the Log2 cluster size of the clusters plotted in b. **d,** c **e,** Barplot showing mean VEGF intensity in GFP+ (mutant) cells from the indicated genotypes normalized by background signal. Data are presented as mean values +/− SD. For statistics see Source Data File 1. Scale bars are 500μm. Abbreviations: P – Postnatal day.

**Extended Data Figure 5:**
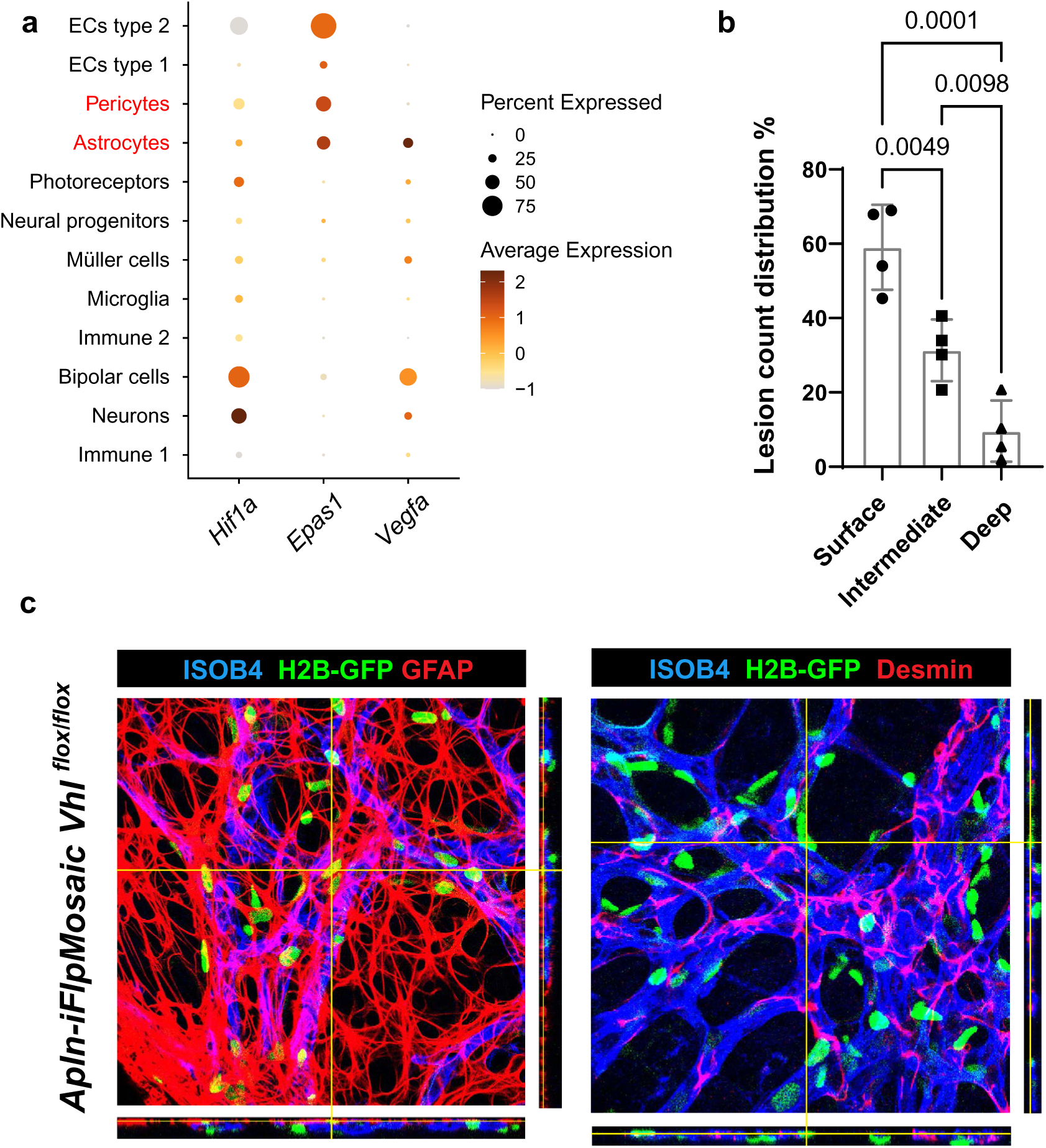
Vhl^KO^ cell clusters in the HB-like lesions show astrocytic features. **a,** Dot plot showing the expression of *Hif1a, Epas1 and Vegfa* in the different postnatal retina clusters. Pericytes and astrocytes are the non-ECs with highest expression of *Epas1.* Among them, astrocytes are the strongest *Vegfa* producers. **b,** Barplot showing how HB-like lession count distribution is mostly localized in the surface layer of the *Apln-iFlpMosaic Vhl^flox/flox^* retina. **c,** Representative 3D confocal micrographs of *Apln-iFlpMosaic Vhl^flox/flox^* HB-like lesions immunostained for ISOB4 and the pericyte Desmin and astrocyte GFAP markers, endogenous GFP detection. Data are presented as mean values +/− SD. For statistics see Source Data File 1. Single cell RNAseq data was reanalysed from Zarkada et.al.

**Extended Data Figure 6:**
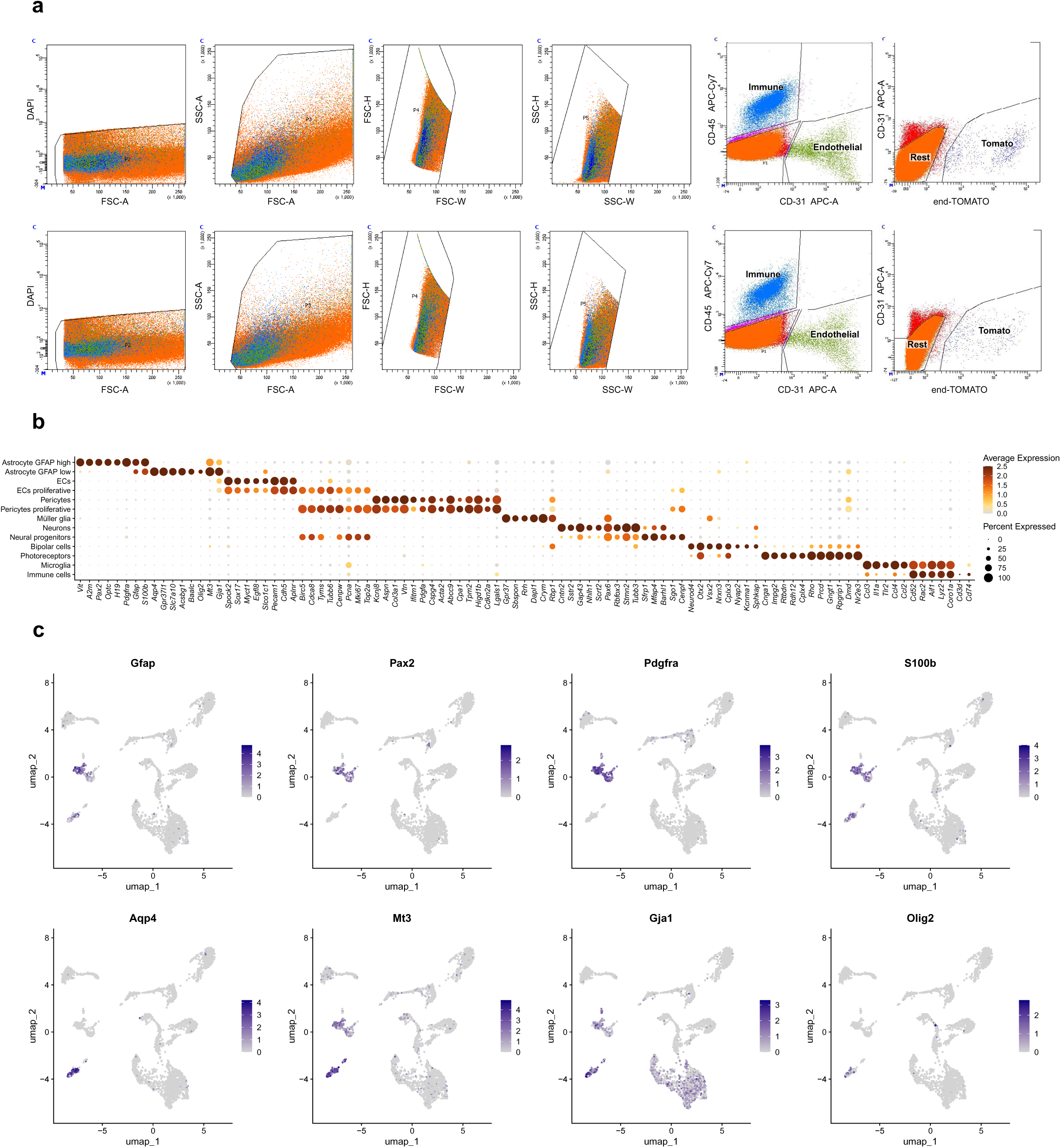
scRNAseq analysis of *GFAP-CreERT2 iSuRe-Cre Vhl* floxed whole retina. **a,** Scatterplots showing the sorting strategy for the scRNAseq analysed cells. Recombined astrocytes (Tomato+), ECs (CD31+ CD45-), Immune cells (CD31-CD45+) and a subset of unlabelled cells were sorted from *GFAP-CreERT2 iSuRe-Cre Vhl^flox/wt^*and *Vhl^flox/flox^* retinas. **b,** Dot plot showing the expression of the different cell type markers. **c,** UMAP showing the expression of markers differentiating astrocyte clusters (GFAP-high and GFAP-low).

**Extended Data Figure 7:**
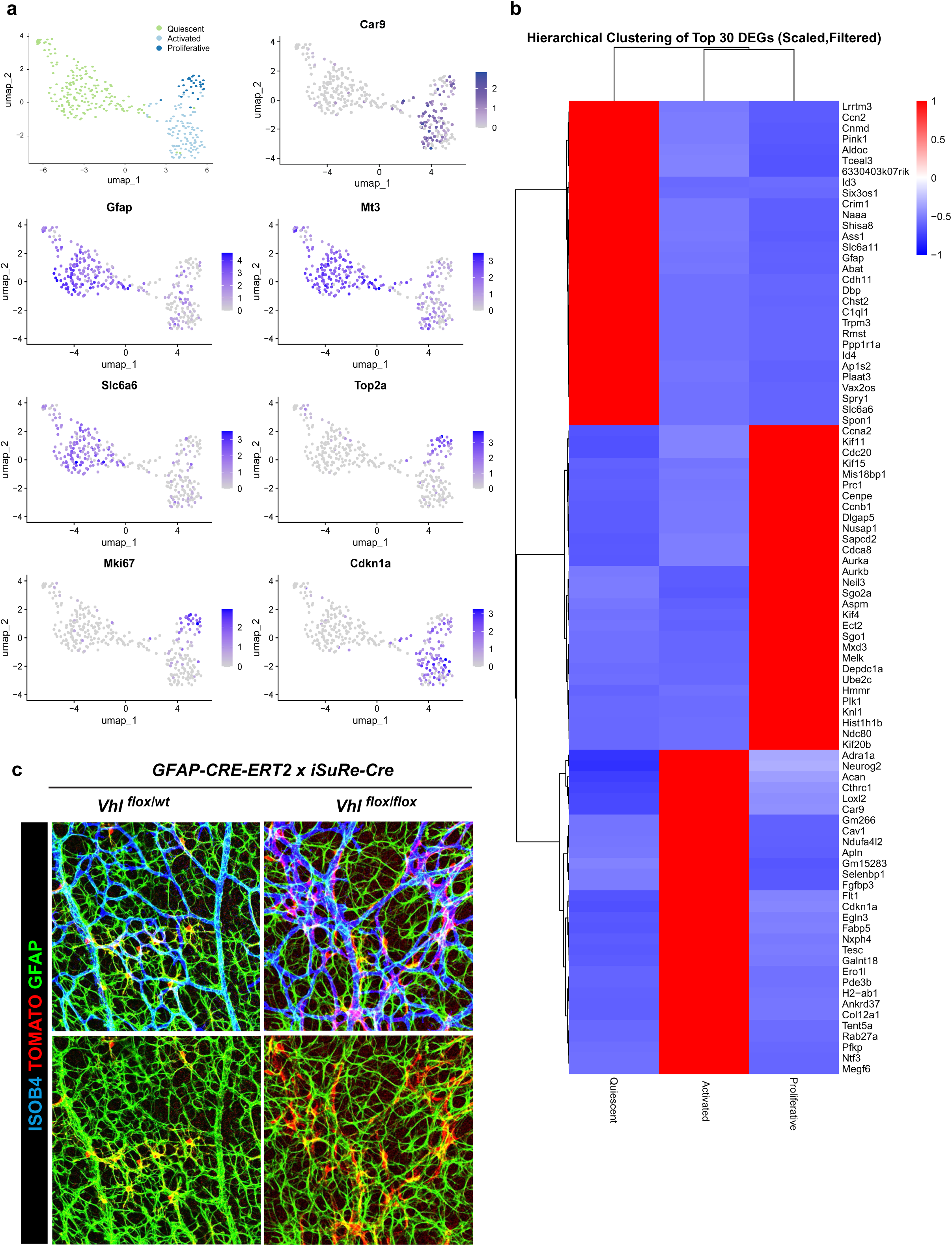
*GFAP-CreERT2 iSuRe-Cre Vhl^flox/flox^*astrocytes show a distinct activated cluster. **a,** UMAP showing the expression of markers differentiating astrocyte clusters (Quiescent, Activated and Proliferative). **b,** Heatmap showing the hierarchical clustering of the top 30 differentially expressed genes (DEGs) for each astrocyte cluster. **c,** Representative confocal micrographs of *GFAP-CreERT2 iSuRe-Cre Vhl^flox/wt^*and *Vhl^flox/flox^* retinas immunostained for ISOB4 and GFAP, endogenous Tomato detection.

**Extended Data Figure 8:**
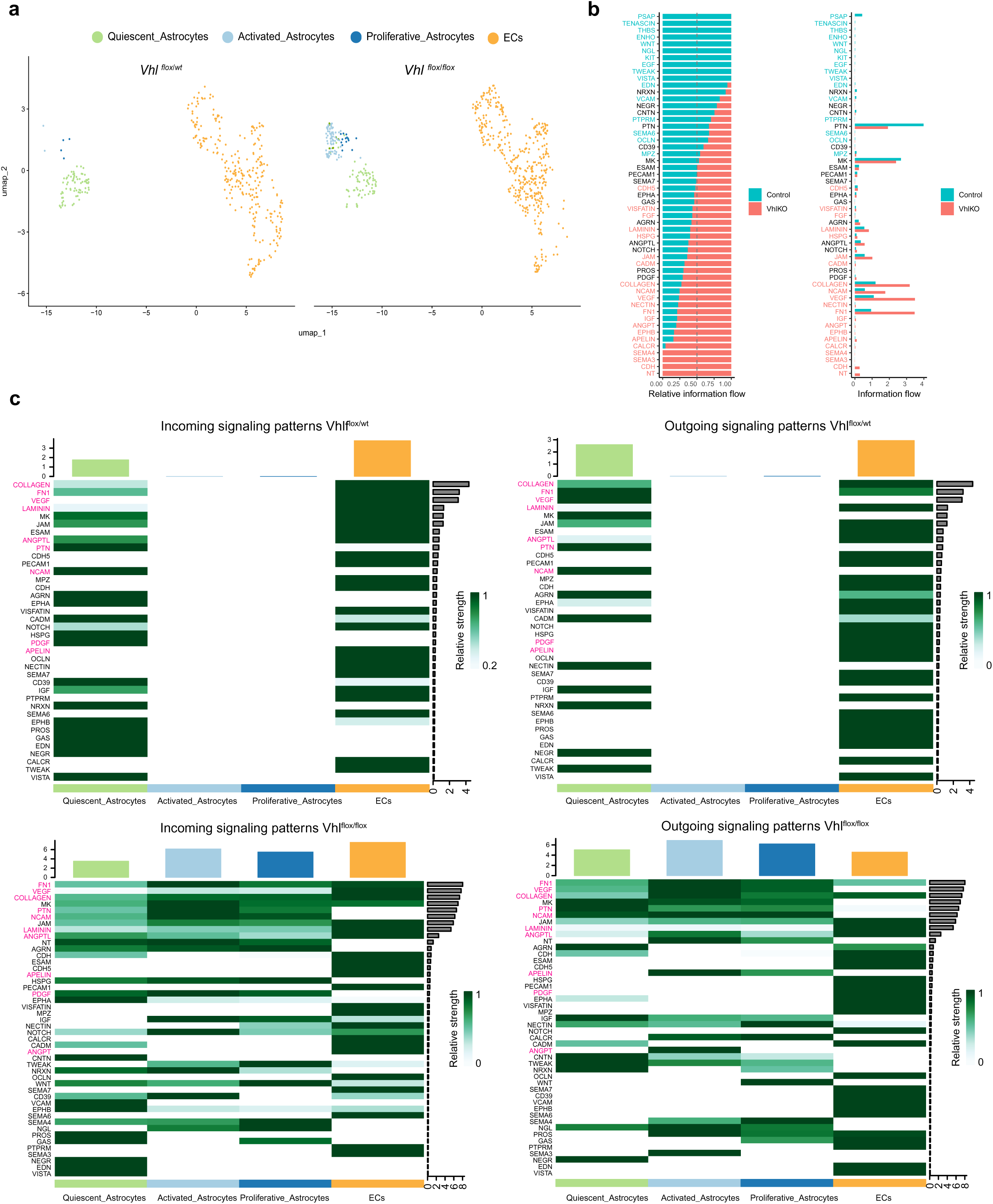
Astrocyte-EC signalling analysis using Cell-Chat. **a,** UMAP showing the clusters that were analysed in the *GFAP-CreERT2, iSuRe-Cre Vhl^flox/wt^* and *Vhl^flox/flox^* conditions. **b,** Full Stacked and Histogram bar graphs representing relative and absolute information flow of the signalling pathways enriched in the *Vhl^flox/wt^* and *Vhl^flox/flox^* conditions. **c,** Heatmap showing the incoming and outgoing signalling patterns representing the pathways with the stronger interactions in both *Vhl^flox/wt^* and *Vhl^flox/flox^*conditions.

**Extended Data Figure 9:**
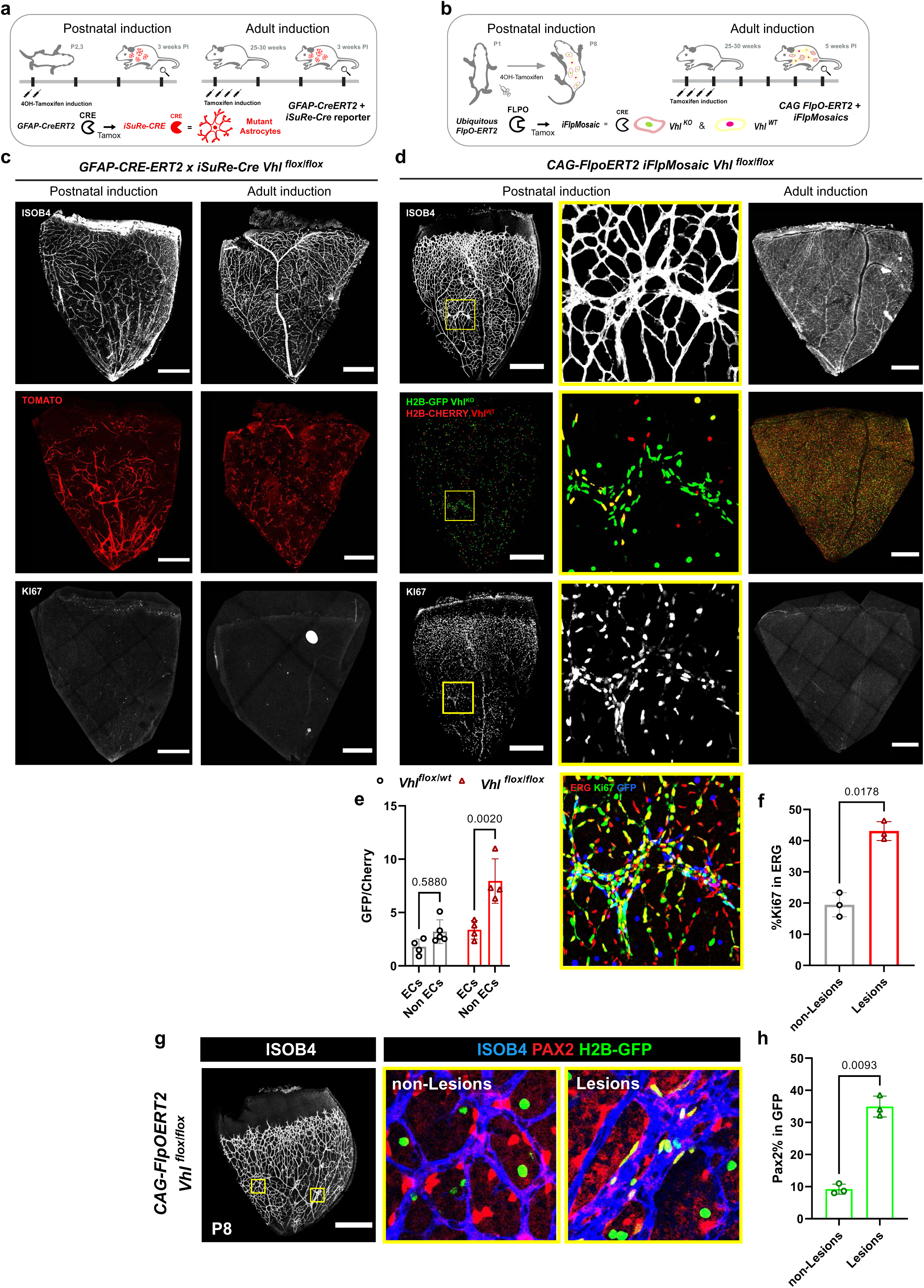
Postnatal, but not adult mosaic deletion of *Vhl* drives HB-like lesions. **a,** Astrocyte specific *GFAP-CreERT2 iSuRe-Cre Vhl^flox/flox^* mice were induced with tamoxifen postnatally or in adulthood. Retinas were dissected 3 weeks after induction. **b,** Ubiquitous CAG-FlpOERT2 iFlpMosaics were postnatally induced with tamoxifen at P1 and analysed at P8 or induced during adulthood and analysed 5 weeks later. **c,** Representative confocal micrographs of postnatal and adult induced *GFAP-CreERT2 iSuRe-Cre Vhl^flox/flox^* retinas immunostained for ISOB4, KI67 and endogenous Tomato detection. HB-like lesions were just observed in the postnatal induced mice. **d,** Representative confocal micrographs of postnatal and adult induced *CAG-FlpOERT2 iFlpMosaic Vhl^flox/flox^* retinas immunostained for ISOB4, KI67 and endogenous GFP and Cherry detection. HB-like lesions are shown in magnified insets; they were just observed in the postnatal induced mice. **e,** Barplot showing the ratio of *Vhl^KO^* (GFP) vs *Vhl^WT^* (Cherry) cells in endothelial and non-endothelial populations in and outside of the HB-like lesions from postnatally induced *CAG-FlpOERT2 iFlpMosaic Vhl^flox/flox^* mice. *Vhl^KO^* GFP+ cells are enriched in the non-EC fraction within the HB-like lesions. **f,** Barplot showing the percentage of KI67+ cells within ECs in and outside of the HB-like lesions in the postnatally induced *CAG-FlpOERT2 iFlpMosaic Vhl^flox/flox^*retinas. **g,** Representative confocal micrographs of postnatally induced *CAG-FlpOERT2 iFlpMosaic Vhl^flox/flox^* retinas immunostained for ISOB4, the astrocyte marker PAX2 and endogenous GFP detection. HB-like lesions are shown in magnified insets. **h,** Barplot showing the percentage of PAX2+ cells within the *Vhl^KO^ GFP* cells in and out of the malformed areas. Data are presented as mean values +/− SD. For statistics see Source Data File 1. Scale bars are 500μm. Abbreviations: P – Postnatal day, VM – Vascular malformation.

**Extended Data Figure 10:**
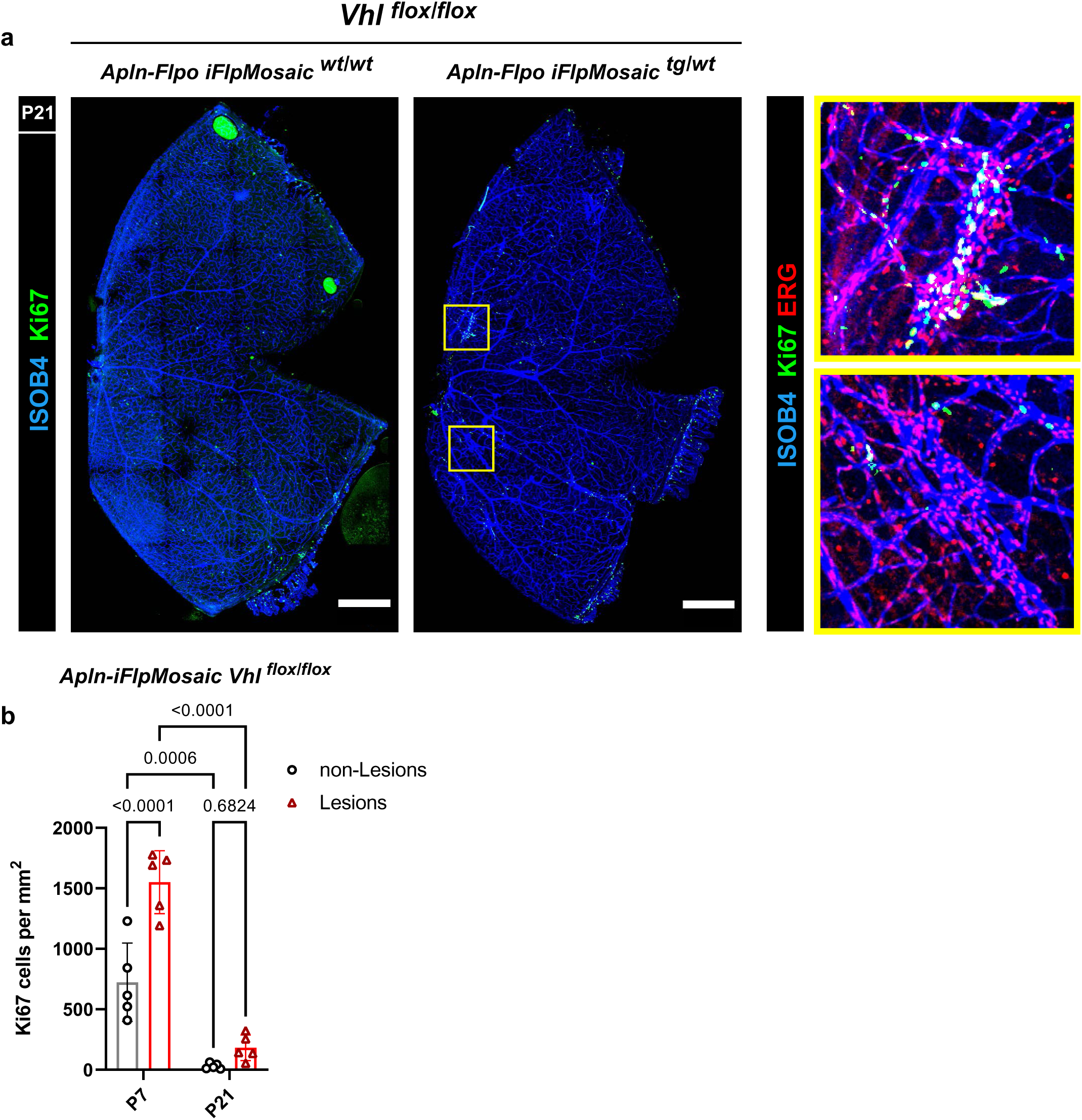
HB-like lesions become mostly quiescent over time. **a,** Representative confocal micrographs of P21 *Apln iFlpMosaic Vhl^flox/flox^* retinas immunostained for ISOB4, KI67 and ERG. KI67+ (Top) and KI67-(Bottom) HB-like lesions are shown in magnified insets. **b,** Barplot showing the percentage of KI67+ cells in capillaries and HB-like lesions from P7 vs P21 retinas show that they both become mostly quiescent by P21. Data are presented as mean values +/− SD. For statistics see Source Data File 1. Scale bars are 500μm. Abbreviations: P – Postnatal day, VM – Vascular malformation.

## Methods

### Mice

All mouse husbandry and experimentation were conducted using protocols approved by local animal ethics committees and authorities (Comunidad Autónoma de Madrid and Universidad Autónoma de Madrid CAM-PROEX 164.8/20, PROEX 293.1/22and PROEX-288.7/24). The mouse colonies were maintained in racked individual ventilation cages according to current national legislation. The mice had dust and pathogen-free bedding and sufficient nesting and environmental enrichment material for the development of species-specific behavior. All mice had ad libitum access to food and water in environmental conditions of 45–65% relative humidity, temperatures of 21–24 °C and a 12–12 h light–dark cycle. To preserve animal welfare, mouse health was monitored with an animal health surveillance program that followed the Federation of European Laboratory Animal Science Associations (FELASA) recommendations for specific pathogen-free facilities

We used Mus musculus lines mostly on the C57BL6 genetic background, with some having a fraction of 129SV, or B6CBAF1 or DBA genetic backgrounds. Mice were backcrossed to C57Bl6 for several generations. To generate mice for analysis, we intercrossed mice aged between 7 and 30 weeks. We do not anticipate any influence on our data of mouse sex. The following mouse lines were used and intercrossed: *Tg(Cdh5-CreERT2)^1Rha^* ^81^*, Tg(GFAP-CreERT2)*^82^*, Tg(INS-CAG-FlpoERT2)*^14^*, Tg(iSuRe-Cre)*^83^*, Vhl^flox^*^16^, *Epas1^flox^*^84^, *Hif1a^flox^*^85^, *Apln-FlpO*^15^, and the *iFlpMosaic* line: *Tg-iFlp^MTomato-2A-H2B-GFP-2A-Cre/MYFP-2A-H2B-Cherry^*^14^.

To induce CreERT2 or FlpO-ERT2 activity in adult mice, 1g tamoxifen (Sigma-Aldrich, P5648_1G) was dissolved in 50 mL corn oil (stock tamoxifen concentration, 20 mg/mL), and aliquots were stored at −20C. Animals received intraperitoneal injections of 100 µL of this stock solution (total dose, 2 mg tamoxifen per animal at 80 mg/kg), as indicated in the figures. To activate recombination in pups, 4-Hydroxytamoxifen (4-OHT) was injected at the indicated stages at a dose of 40 mg/kg. All mouse lines and primer sequences required for genotyping are provided in **Supplementary Table 1**.

### Immunostaining of whole retinas

For immunostaining of mouse retinas, eyes were dissected from mouse pups and fixed by incubation with agitation for 20 min in 4% PFA in PBS (diluted from a stock of 16% PFA; EMS 15710). After two washes in PBS, retinas were microdissected from the eyes, 4 incisions made to enable their flat mount later, and refixed in 4% PFA for 45 min with agitation. Dissected retinas were blocked and permeabilized by incubation for 1 h in PBST (0.3% Triton X-100, 3% fetal bovine serum (FBS) and 3% donkey serum in PBS). Samples were then washed twice in PBST and incubated with primary antibodies (**Supplementary Table 2**) diluted in PBST overnight at 4°C with agitation. The next day, retinas were washed 5 × 20 min in PBST diluted 1:2. And after incubated in this same solution for 2 h at room temperature with Alexa-conjugated secondary antibodies (**Supplementary Table 2**). After 3 × 15 min washes in PBS containing 0.15% Triton X-100, retinas were washed 2 × 15 min in PBS and flat mounted in Fluoromount-G (SouthernBiotech).

### Image acquisition and analysis

For confocal scanning, the immunostained whole-mount retinas were imaged at high resolution with a Leica SP8 Navigator confocal microscope fitted with a 20× or 40× objective. Individual fields or tiles of large areas were acquired. All the images shown are representative of the results obtained for each group and experiment. The animals were dissected and processed under exactly the same conditions. Comparisons of phenotypes or signal intensities were made using images obtained with the same laser excitation and confocal scanner detection settings. ImageJ/FIJI v1.54f was used to threshold, select and quantify the objects in confocal micrographs.

Quantification of mouse retina confocal images was performed using ImageJ/Fiji software. We developed and adapted several macros in IJ1Macro language to enable fully or semi-automated analysis. Thresholds were either automatically calculated or manually set depending on signal intensity and signal-to-noise ratio. Core plugins used included Watershed (Soille and Vincent, 1990), Analyse Particles, Spatial Statistics, Bio-Formats, TrackMate, and other pre-installed tools. Particle identification was carried out by segmenting nuclear signals using the Analyse Particles plugin. Identified regions of interest (ROIs) were then interrogated for signal in other channels of interest and classified as positive or negative based on defined thresholds.

Endothelial cell (EC) density was calculated as the number of ERG⁺ cells per plexus area. Frequencies of Ki67⁺ in ECs or reporters (GFP, Cherry or Tomato) were semi-automatically quantified as the proportion of double-positive cells relative to the total number of cells per field. In *iFlpMosaic* retinas, GFP and Cherry cell ratios were quantified by independent segmentation and counting of nuclear signals. VEGF and P21 intensity within defined areas (Tomato⁺, IB4⁺, or GFP^+^) was quantified by calculating the mean VEGF signal within the thresholded region and normalized to background signal from the VEGF channel in each image.

For clonal analysis, the X-Y coordinates of all GFP⁺ and Cherry⁺ particles were extracted and exported. Distance measurements between cells expressing the same reporter type were performed using an R script in which cells located within 20μm of each other were considered part of the same clone. Clonal size and frequency distributions were plotted as histograms using GraphPad Prism.

### Latex perfusion

For latex perfusion, pink latex (provided by Mariona Graupera, Josep Carreras Leukaemia Research Institute) was diluted in PBS at a 9:1 (latex:PBS) ratio to facilitate injection. Adult mice were euthanized in a CO₂ chamber and the chest was opened to expose the heart. After puncturing the right atrium to open the circulatory circuit, 10 ml of PBS was perfused through the left ventricle to flush out the blood. The abdominal cavity was opened carefully, avoiding damage to blood vessels, to monitor the effectiveness of the washout. Once blood was cleared, the diluted latex was slowly perfused through the same left ventricle (using the same entry point if possible). The skin of the right leg was removed to visualize the femoral artery, and perfusion was stopped when latex reached the right femoral artery at the quadriceps. After waiting 5 minutes to allow the latex to solidify and prevent leakage, organs were dissected, rinsed in PBS, and fixed overnight in 4% paraformaldehyde (PFA) at 4 °C. The next day, organs were washed in PBS three times for 10 minutes each at 4 °C.

### CUBIC-1 clearing of latex perfused organs

To achieve optical clearing of latex perfused mouse organs, adult fixed organs were incubated in freshly prepared CUBIC Reagent-1, composed of 25% urea, 25% N,N,N’,N’-tetrakis(2-hydroxypropyl) ethylenediamine, 25% Triton X-100, and 35% distilled water (dH₂O), for 2 days at 37 °C to remove lipids. The reagent was refreshed once after the first 24 hours of incubation. Organs were subsequently imaged in CUBIC-1 using a Leica dissecting microscope.

### Lectin perfusion and EZ-Clear clearing

Adult mice (P21) were anesthetized with an intraperitoneal injection of 100 µL pentobarbital mixed with lidocaine. Under deep anesthesia, mice were retro-orbitally injected using a 31-gauge insulin syringe (BD, 328438) with 50 µL of far-red (649 nm) fluorescently conjugated lectin (Lycopersicon Esculentum Lectin, DyLight 649, Invitrogen L32472) to label the luminal endothelium of patent vessels. The lectin was allowed to circulate for 5 minutes. Following circulation, the thoracic cavity was opened, the heart was exposed, and the right atrium was incised to allow drainage. Mice were then perfused with 10 mL of phosphate-buffered saline (PBS), followed by 10 mL of 4% paraformaldehyde (PFA).

Brain was carefully dissected and post-fixed in 4% PFA at 4°C for 24 hours, and washed 3 times for 15 min with PBS. Organs were then placed in individual glass scintillation vials and wrapped in aluminum foil to protect them from light. Samples were incubated in 20 mL of a 50% (v/v) tetrahydrofuran/Milli-Q water solution for 16 hours on an orbital shaker inside a vented chemical hood. Following incubation, samples were rinsed four times with 20 mL of sterile Milli-Q water for 1 hour at room temperature. After washing, samples were incubated in 5 mL of EZ View solution (80% Nycodenz, 7 M urea, and 0.05% sodium azide in 0.02 M sodium phosphate buffer) for 24 hours. Samples were then visualized under a Leica SP8 Navigator confocal microscope (10X objective) while submerged in EZ View buffer.

### In vivo pharmacological treatments

For the VEGF blocking treatment we intraperitoneally injected mice at the indicated postnatal stages with Aflibercept (Zaltrap, 25mg/kg) diluted in PBS.

For the mTORC1 inhibition treatment we intraperitoneally injected mice at the indicated postnatal stages with rapamycin at a final concentration of 4mg/kg. Prior to administration, stock rapamycin solution (25mg/mL dissolved in DMSO) was diluted in a vehicle solution composed by PEG300 (30.3%), Tween-80 (5.1%), and phosphate-buffered saline (PBS; 64.6%).

### Analysis of publicly available single cell data

As described in the main text, single-cell RNA-sequencing data from wild-type (WT) P6 mouse retinas were obtained from Zarkada et al. (2021)^22^. Raw count matrices were imported using the Read10X function and processed into a Seurat object via the CreateSeuratObject function. Low-quality cells were filtered out based on standard quality control criteria, including low gene counts and high mitochondrial gene content (percentage of reads mapping to mitochondrial genes).

Quality metrics—including gene count per cell, total UMI count, and mitochondrial content—were visualized using violin plots. Data normalization was performed using the NormalizeData function with log transformation and a fixed scaling factor. Highly variable genes were identified with the FindVariableFeatures function using the “vst” selection method. All genes were scaled prior to dimensionality reduction to mitigate technical variation.

Principal component analysis (PCA) was conducted using the RunPCA function. The number of principal components retained for downstream analysis was determined using JackStraw analysis and elbow plots. Cell clustering was performed using the FindNeighbors and FindClusters functions (Louvain algorithm), with tuning of the resolution parameter to optimize cluster number.

Cluster-specific marker genes were identified using the FindMarkers function, considering only significantly upregulated (positive) markers in each cluster. Based on expression profiles of canonical markers and literature data, clusters were annotated with putative retinal cell type identities. These identities were incorporated into the Seurat object metadata.

Gene expression across clusters was visualized using the DotPlot function for selected markers of interest. The fully processed Seurat object, including clustering and cell-type annotations, was saved as an .RDS file for downstream analyses.

### Cell isolation for transcriptomic analysis

P8 retinas were freshly dissected and dissociated using the Neural Tissue Dissociation Kit – Postnatal Neurons from Miltenyi Biotec (CAT number: 130-094-802).

For RNA-seq analysis, dissociated retinas were passed through a 70 μm cell strainer and centrifuged (450g, 4 °C, 5 min). Cells were resuspended in antibody incubation solution (PBS without Ca²⁺ or Mg²⁺, supplemented with 2% dialyzed FBS; Biowest X0515) together with primary antibodies: APC rat anti-mouse CD31 (labels ECs), APC-Cy7 rat anti-CD45 (labels blood cells) and HTO-antibodies (BD Biosciences, **Supplementary Table 2,** to hashtag and distinguish the different genotypes when loaded on the same 10x Genomics port. Samples were then incubated at 4 °C with gentle agitation for 20 minutes. After antibody incubation, samples were washed twice with antibody incubation solution, centrifuged (450 × *g*, 5 min, 4 °C), and resuspended in sorting buffer (10% FBS in PBS without Ca²⁺ or Mg²⁺). Stained cells were maintained on ice until acquisition. Immediately prior to FACS analysis, DAPI (5 mg ml⁻¹) was added and the cell suspension was passed slowly through a 70 µm cell strainer attached to a 1 ml syringe to remove aggregates.

Alive, DAPI-negative, CD31+/CD45-, CD31-/CD45+, endogenous reporter positive and negative cells were sorted into cold 1.5 ml microcentrifuge tubes containing 300 µl of sorting buffer. Cells were collected using a 100 µm nozzle at 20 PSI under high-purity mode and a flow rate below 3,000 events per second to preserve cell viability and minimize contamination. After sorting, cells were pelleted at 500g for 5 minutes at 4C, and resuspended in 30–40 µl cell-capture buffer (Ca^2+^ and Mg^2+^ free PBS supplemented with 0.04% ultra-pure BSA, Thermofisher AM2616). Viability and cell concentration were determined using a Countess 3 Automated Cell Counter (Thermo Fisher Scientific) before loading the 10x Genomics port.

### Next-generation sequencing sample and library preparation

Single-cell suspensions were loaded onto the 10x genomics Chromium Controller system (10x Genomics) for encapsulation into emulsion droplets. The 10x genomics Chromium single cell 3’ RNA-seq library preparation and sequencing were carried out at the CNIC Genomics Unit according to the manufacturer’s protocols. The port was loaded with 30k cells. Final libraries were sequenced on an Illumina NextSeq 2000 platforms. Sequencing was performed with an average depth of 41,831 reads/cell.

### Single cell RNA data analysis

Single-cell RNA-seq data were aligned and quality controlled by the CNIC Bioinformatics Unit. Clustering, cell type identification and downstream analysis was performed by Macarena de Andrés. The following pipeline was applied for transcript alignment, quantification, quality control, and cell-type classification.

#### Pre-processing and alignment

Reference transcriptomes were built using the GRCm38 mouse genome and Ensembl gene build v98 (September 2019, sep2019.archive.ensembl.org). Gene annotations were obtained from the corresponding Ensembl BioMart archive. Raw sequencing reads were aligned and quantified using Cell Ranger v7.1.0.

#### Cell filtering, demultiplexing, and normalization

Downstream analysis was performed using Seurat v4.1.3. Cells were filtered using the following criteria: total UMI counts between 1,500 and 60,000; detection of more than 800 genes per cell; mitochondrial transcript content less than 25%; ribosomal transcript content was assessed but not used for filtering; cells contributing at least 0.2% of total UMIs in the sample; less than 65% of counts attributed to the top 50 expressed genes; and haemoglobin gene (HBB) content below 0.1%. Additionally, cells with fewer than 25 hashtag oligonucleotide (HTO) UMIs were removed to improve sample demultiplexing. Cells were demultiplexed using sample-specific hashtag antibody signals (**Supplementary Table 2**).

#### Clustering, doublet removal and cell type identification

Cells were clustered by constructing a shared nearest neighbour (SNN) graph and applying the Louvain community detection algorithm to identify distinct cell populations. The optimal number of principal components for dimensionality reduction was determined using an Elbow plot, and clustering resolution parameters were selected based on the sample complexity.

Putative doublets were identified as cell clusters expressing high total UMI/gene counts, unusual gene co-expression and located in PCA outliers.

Final cluster identities were manually curated based on literature-based cell markers.

#### Cell-cell communication analysis

Annotated astrocyte clusters (quiescent, activated, and proliferative) and endothelial cell (EC) clusters were merged into a single Seurat object using the merge function. The combined dataset was normalized, and variable features were identified prior to scaling. Principal component analysis (PCA) was performed to reduce dimensionality, followed by determination of significant principal components using JackStraw analysis and elbow plots. Cells were clustered based on the first 20 PCs at a resolution of 0.5 and visualized using UMAP.

To simplify downstream analysis, endothelial clusters were merged into a single EC cluster. The resulting object was saved for further analysis. Subsets corresponding to *Vhl^flox/flox^* and *Vhl^flox/wt^*genotypes were generated. Cell-cell communication networks were inferred separately for *Vhl^flox/flox^* and *Vhl^flox/wt^* datasets using the CellChat^61,62^ R package with the mouse CellChat database restricted to secreted signalling, cell-cell contact, and ECM-receptor interactions. Overexpressed genes and interactions were identified, and communication probabilities were computed with a trimean method. Low-confidence interactions (fewer than 10 cells) were filtered out before pathway-level communication probabilities were aggregated. Network centrality analyses were performed to determine signalling roles across clusters. Differential communication patterns between *Vhl^flox/flox^* and *Vhl^flox/wt^*conditions were visualized via circle plots, heatmaps, and bubble plots. Finally, merged CellChat objects enabled comparative analyses of interaction counts, strengths, and information flow to identify condition-specific changes in ligand-receptor signalling.

### Micro Computed Tomography

CT studies were performed with a small-animal PET/CT scanner (nanoScan, Mediso, Hungary). During the acquisition, mice were anesthetized using isoflurane 2% and 1.8 L/min oxygen flow. Ophthalmic gel was placed in the eyes to prevent drying.

CT images were acquired 30-60 minutes after intravenous administration of 0.1 ml of 2000 mg/ml of gold nanoparticle (AuNPs) using an X-ray beam current of 178 µA, a tube voltage of 45 kVp and 500 ms of exposure. CT data was reconstructed using a Ramlack algorithm with a voxel dimension of 0.14 mm3. A 3D Maximum Intensity Projection (MIP) of the whole body was obtained using the Multimodality Workstation software.

AuNPs were synthesized using a modified Turkevich method. This classical approach involves the reduction of tetrachloroauric acid (HAuCl₄) by trisodium citrate, which also acts as a stabilizing agent. In a typical synthesis, 1 liter of 0.5 mM HAuCl₄ solution was prepared in a round-bottom flask. Separately, 100 mL of 50 mM trisodium citrate solution was preheated to 70 °C. The gold precursor solution was brought to a vigorous boil under constant stirring using a heating mantle. Once the boiling point was reached, the preheated trisodium citrate solution was rapidly poured onto the boiling HAuCl₄ solution. The reaction mixture was maintained under boiling and continuous stirring for 30 minutes, during which a distinct color change—from pale yellow to deep red—indicated the formation of colloidal gold nanoparticles. When the color stabilized, the solution was allowed to cool naturally to room temperature. To remove residual reactants and byproducts, the AuNP solution was dialyzed against 5 mM sodium citrate using a dialysis membrane with a molecular weight cut-off (MWCO) of 10,000 Da. Following synthesis and purification, the AuNPs were subjected to a surface functionalization (pegylation) process to enhance biocompatibility and colloidal stability. First, 1 mL of 1 mM methoxy polyethylene glycol thiol (mPEG-SH, Mw = 5,000 g/mol) was added to the nanoparticle solution, which was then stirred at room temperature for 1 hour. Subsequently, 1 mL of 2 mM mPEG-SH (Mw = 1,000 g/mol) was introduced to the mixture. The total volume was concentrated by rotatory evaporation to 10 mL and the solution was dialyzed with miliQ water to remove unbound PEG molecules.

Finally, the nanoparticle suspension was concentrated by rotary evaporation to a final volume of 0.5 mL, resulting in a gold nanoparticle dispersion with a concentration of approximately 200 mg/mL. This PEGylated AuNP formulation (AuCt) is suitable for use as a computed tomography (CT) contrast agent due to its high electron density, colloidal stability, and biocompatibility.

### Statistics and reproducibility

Most numerical data shown in charts was first compiled and processed with Microsoft Excel 2019 and after analysed and plotted with Graphpad Prism v10.1.0. All bar graphs show mean ± standard deviation. The experiments were repeated with independent animals, as stated in the source data file or figure legends. The comparisons between two sample groups with a Gaussian distribution were by unpaired two-tailed Student t-tests. The comparisons among more than two groups were done by one-way or 2-way analysis of variance followed by multiple comparison tests. Datapoints were analysed and plotted with GraphPad Prism. Images in which particle selection was performed manually were blinded. Animals or tissues were selected for analysis based on their genotype, the detected Cre/FlpO-dependent recombination frequency and the quality of multiplex immunostaining. The sample sizes were chosen according to the observed statistical variation.

## References

1. Conway, J. E. et al. Hemangioblastomas of the Central Nervous System in von Hippel-Lindau Syndrome and Sporadic Disease. Neurosurgery 48, 55–63 (2001).

2. Gläsker, S., Vergauwen, E., Koch, C. A., Kutikov, A. & Vortmeyer, A. O. Von Hippel-Lindau Disease : Current Challenges and Future Prospects. 5669–5690 (2020).

3. Larcher, A. et al. Multidisciplinary management of patients diagnosed with von Hippel-Lindau disease: A practical review of the literature for clinicians. Asian Journal of Urology 9, 430–442 (2022).

4. Iliopoulos, O. et al. Belzutifan for patients with von Hippel-Lindau disease-associated CNS haemangioblastomas (LITESPARK-004): a multicentre, single-arm, phase 2 study. The Lancet Oncology 25, 1325–1336 (2024).

5. Shankar, G. M. et al. Sporadic hemangioblastomas are characterized by cryptic VHL inactivation. Acta Neuropathologica Communications 2, (2014).

6. Vortmeyer, A. O. et al. von Hippel-Lindau gene deletion detected in the stromal cell component of a cerebellar hemangioblastoma associated with von Hippel-Lindau disease. Human Pathology 28, 540–543 (1997).

7. Shively, S. B. et al. Developmentally arrested structures preceding cerebellar tumors in von Hippel-Lindau disease. Modern Pathology 24, 1023–1030 (2011).

8. Park, D. M. et al. Von Hippel-Lindau disease-associated hemangioblastomas are derived from embryologic multipotent cells. PLoS Medicine 4, 0333–0341 (2007).

9. Pilotto, E. et al. Retinal glial cells in von hippel–lindau disease: A novel approach in the pathophysiology of retinal hemangioblastoma. Cancers 14, 1–14 (2022).

10. Miyazawa, A. et al. Expression of inhibin α by stromal cells of retinal angiomas excised from a patient with von Hippel-Lindau disease. Japanese Journal of Ophthalmology 53, 501–505 (2009).

11. Wang, H. et al. Deletion of the von Hippel-Lindau gene in hemangioblasts causes Hemangioblastoma-like Lesions in Murine Retina. Cancer Research 78, 1266–1274 (2018).

12. Wei, R. et al. Rb1/Rbl1/Vhl loss induces mouse subretinal angiomatous proliferation and hemangioblastoma. JCI Insight 4, 1–19 (2019).

13. AO Vortmeyer et al. Evolution of VHL tumourigenesis in nerve root tissue. 231–241 (2006) doi:10.1002/path.

14. Garcia-Gonzalez, I. et al. iFlpMosaics enable the multispectral barcoding and high-throughput comparative analysis of mutant and wild-type cells. Nat Methods 22, 323–334 (2025).

15. Luo, W. et al. Arterialization requires the timely suppression of cell growth. Nature 589, 437–441 (2021).

16. Haase, V. H., Glickman, J. N., Socolovsky, M. & Jaenisch, R. Vascular tumors in livers with targeted inactivation of the von Hippel–Lindau tumor suppressor. Proceedings of the National Academy of Sciences 98, 1583–1588 (2001).

17. Otero-Marquez, O. et al. 3-D OCT angiographic evidence of Anti-VEGF therapeutic effects on retinal capillary hemangioma. American Journal of Ophthalmology Case Reports 25, 101394 (2022).

18. Chew, M. Y. Ocular manifestations of von Hippel-Lindau disease: Clinical and genetic investigations. Transactions of the American Ophthalmological Society 103, 495–511 (2005).

19. Ola, R. et al. PI3 kinase inhibition improves vascular malformations in mouse models of hereditary haemorrhagic telangiectasia. Nature Communications 7, (2016).

20. Klingler, J.-H. et al. Hemangioblastoma and von Hippel-Lindau disease: genetic background, spectrum of disease, and neurosurgical treatment. Child’s Nervous System (2020) doi:10.1007/s00381-020-04712-5/Published.

21. Fernandes, D. A. et al. Imaging manifestations of von Hippel-Lindau disease: an illustrated guide focusing on abdominal manifestations. Radiologia Brasileira 55, 317–323 (2022).

22. Zarkada, G. et al. Specialized endothelial tip cells guide neuroretina vascularization and blood-retina-barrier formation. Developmental Cell 56, 2237–2251.e6 (2021).

23. Leung David W., Cachianes George, Kuang Wun-Jing, Goeddel David V., & Ferrara Napoleone. Vascular endothelial growth factor is a secreted angiogenic mitogen. Science 246, 1306–1309 (1989).

24. Levy, A. P., Levy, N. S. & Goldberg, M. A. Hypoxia-inducible protein binding to vascular endothelial growth factor mRNA and its modulation by the von Hippel-Lindau protein. Journal of Biological Chemistry 271, 25492–25497 (1996).

25. Rattner, A., Williams, J. & Nathans, J. Roles of HIFs and VEGF in angiogenesis in the retina and brain. Journal of Clinical Investigation 129, 3807–3820 (2019).

26. Chan, Chi-Chao, M., et al. VHL Gene Deletion and Enhanced VEGF Gene Expression Detected in the Stromal Cells of Retinal Angioma. 117, 625–630 (1999).

27. Vortmeyer, A. O. et al. von Hippel-Lindau gene deletion detected in the stromal cell component of a cerebellar hemangioblastoma associated with von Hippel-Lindau disease. Human Pathology 28, 540–543 (1997).

28. Gerhardt, H. et al. VEGF guides angiogenic sprouting utilizing endothelial tip cell filopodia. Journal of Cell Biology 161, 1163–1177 (2003).

29. Semenza, G. L. Oxygen sensing, hypoxia-inducible factors, and disease pathophysiology. Annual Review of Pathology: Mechanisms of Disease 9, 47–71 (2014).

30. Peres, T., Aeppli, S., Fischer, S., Hundsberger, T. & Rothermundt, C. First Single-Centre Experience with the Novel HIF-α Inhibitor Belzutifan in Switzerland. Curr Oncol 32, 64 (2025).

31. Jonasch, E. et al. Belzutifan for Renal Cell Carcinoma in von Hippel–Lindau Disease. New England Journal of Medicine 385, 2036–2046 (2021).

32. Stone, J. et al. Development of retinal vasculature is mediated by hypoxia-induced vascular endothelial growth factor (VEGF) expression by neuroglia. Journal of Neuroscience 15, 4738– 4747 (1995).

33. Paisley, C. E. & Kay, J. N. Seeing stars: Development and function of retinal astrocytes. Developmental Biology 478, 144–154 (2021).

34. donato2001.

35. Eng, L. F., Vanderhaeghen, J. J., Bignami, A. & Gerstl, B. An acidic protein isolated from fibrous astrocytes. Brain Research 28, 351–354 (1971).

36. Delpech, B. et al. Glial fibrillary acidic protein in tumours of the nervous system. Br J Cancer 37, 33–40 (1978).

37. Bosze, B. et al. Multiple roles for *Pax2* in the embryonic mouse eye. Developmental Biology 472, 18–29 (2021).

38. Tatsumi, K. et al. Olig2-Lineage Astrocytes: A Distinct Subtype of Astrocytes That Differs from GFAP Astrocytes. Front. Neuroanat. 12, (2018).

39. Paisley, C. E. & Kay, J. N. Seeing stars: Development and function of retinal astrocytes. Developmental Biology 478, 144–154 (2021).

40. Liddelow, S. A. et al. Neurotoxic reactive astrocytes are induced by activated microglia. Nature 541, 481–487 (2017).

41. Del Peso, L. et al. The von Hippel Lindau/Hypoxia-inducible Factor (HIF) Pathway Regulates the Transcription of the HIF-Proline Hydroxylase Genes in Response to Low Oxygen. Journal of Biological Chemistry 278, 48690–48695 (2003).

42. Lai, R. K.-H. et al. NDUFA4L2 Fine-tunes Oxidative Stress in Hepatocellular Carcinoma. Clinical Cancer Research 22, 3105–3117 (2016).

43. Mesa-Ciller, C. et al. Unique expression of the atypical mitochondrial subunit NDUFA4L2 in cerebral pericytes fine tunes HIF activity in response to hypoxia. J Cereb Blood Flow Metab 43, 44–58 (2023).

44. Fukuda, R. et al. HIF-1 Regulates Cytochrome Oxidase Subunits to Optimize Efficiency of Respiration in Hypoxic Cells. Cell 129, 111–122 (2007).

45. Chatzopoulos, K., Aubry, M.-C. & Gupta, S. Immunohistochemical expression of carbonic anhydrase 9, glucose transporter 1, and paired box 8 in von Hippel-Lindau disease–related lesions. Human Pathology 123, 93–101 (2022).

46. Laird, M. et al. Mitochondrial metabolism regulation and epigenetics in hypoxia. Front. Physiol. 15, (2024).

47. Bao, M. H. R. & Wong, C. C. L. Hypoxia, metabolic reprogramming, and drug resistance in liver cancer. Cells 10, 1–18 (2021).

48. Corsetto, P. et al. Effects of Germline VHL Deficiency on Growth, Metabolism, and Mitochondria. 835–844 (2020) doi:10.1056/NEJMoa1907362.

49. Cao, Y., Langer, R. & Ferrara, N. Targeting angiogenesis in oncology, ophthalmology and beyond. Nat Rev Drug Discov 22, 476–495 (2023).

50. Helker, C. S. M. et al. Apelin signaling drives vascular endothelial cells towards a pro-angiogenic state. eLife 9, 1–44 (2020).

51. Reinhard, J., Wiemann, S., Hildebrandt, S. & Faissner, A. Extracellular Matrix Remodeling in the Retina and Optic Nerve of a Novel Glaucoma Mouse Model. Biology (Basel) 10, 169 (2021).

52. Tao, C. & Zhang, X. Retinal Proteoglycans Act as Cellular Receptors for Basement Membrane Assembly to Control Astrocyte Migration and Angiogenesis. Cell Reports 17, 1832–1844 (2016).

53. Hara, M. et al. Interaction of reactive astrocytes with type I collagen induces astrocytic scar formation through the integrin–N-cadherin pathway after spinal cord injury. Nat Med 23, 818– 828 (2017).

54. Puebla, M., Tapia, P. J. & Espinoza, H. Key Role of Astrocytes in Postnatal Brain and Retinal Angiogenesis. International Journal of Molecular Sciences 23, (2022).

55. Tao, C. & Zhang, X. Development of astrocytes in the vertebrate eye. Developmental Dynamics 243, 1501–1510 (2014).

56. White, R. E. et al. Transforming Growth Factor α Transforms Astrocytes to a Growth-Supportive Phenotype after Spinal Cord Injury. J. Neurosci. 31, 15173–15187 (2011).

57. White, R. E., Yin, F. Q. & Jakeman, L. B. TGF-alpha increases astrocyte invasion and promotes axonal growth into the lesion following spinal cord injury in mice. Exp Neurol 214, 10–24 (2008).

58. Bohrer, C. et al. The balance of Id3 and E47 determines neural stem/precursor cell differentiation into astrocytes. EMBO J 34, 2804–2819 (2015).

59. Wanner, I. B. et al. Glial Scar Borders Are Formed by Newly Proliferated, Elongated Astrocytes That Interact to Corral Inflammatory and Fibrotic Cells via STAT3-Dependent Mechanisms after Spinal Cord Injury. J. Neurosci. 33, 12870–12886 (2013).

60. Verkhratsky, A. et al. Astrocytes in human central nervous system diseases: a frontier for new therapies. Sig Transduct Target Ther 8, 1–37 (2023).

61. Jin, S. et al. Inference and analysis of cell-cell communication using CellChat. Nat Commun 12, 1088 (2021).

62. Jin, S., Plikus, M. V. & Nie, Q. CellChat for systematic analysis of cell–cell communication from single-cell transcriptomics. Nat Protoc 20, 180–219 (2025).

63. Biswas, S., Bachay, G., Chu, J., Hunter, D. D. & Brunken, W. J. Laminin-Dependent Interaction between Astrocytes and Microglia: A Role in Retinal Angiogenesis. The American Journal of Pathology 187, 2112–2127 (2017).

64. Fruttiger, M. et al. PDGF mediates a neuron-astrocyte interaction in the developing retina. Neuron 17, 1117–1131 (1996).

65. Gerhardt, H. et al. VEGF guides angiogenic sprouting utilizing endothelial tip cell filopodia. Journal of Cell Biology 161, 1163–1177 (2003).

66. Håkansson, J., Ståhlberg, A., Sand, F. W., Gerhardt, H. & Semb, H. N-CAM Exhibits a Regulatory Function in Pathological Angiogenesis in Oxygen Induced Retinopathy. PLOS ONE 6, e26026 (2011).

67. Mansur, A. & Radovanovic, I. Vascular malformations: An overview of their molecular pathways, detection of mutational profiles and subsequent targets for drug therapy. Front. Neurol. 14, (2023).

68. Nisbet, R. et al. Oral sirolimus therapy for patients with complex low-flow vascular malformations. Journal of Vascular Surgery: Venous and Lymphatic Disorders 102261 (2025) doi:10.1016/j.jvsv.2025.102261.

69. Cavazos, R. et al. Sirolimus for vascular anomalies in the first year of life: a systematic review. J Perinatol 44, 1087–1097 (2024).

70. Park, S. & Chan, C.-C. Von Hippel-Lindau Disease (VHL): A need for a murine model with retinal hemangioblastoma. Histol Histopathol. 27, 975–984 (2012).

71. Weidemann, A. et al. Astrocyte hypoxic response is essential for pathological but not developmental angiogenesis of the retina. Glia 58, 1177–1185 (2010).

72. Kurlekar, S. et al. Oncogenic Cell Tagging and Single-Cell Transcriptomics Reveal Cell Type– Specific and Time-Resolved Responses to *Vhl* Inactivation in the Kidney. Cancer Research 84, 1799–1816 (2024).

73. Gläsker, S. et al. Hemangioblastomas share protein expression with embryonal hemangioblast progenitor cell. Cancer Research 66, 4167–4172 (2006).

74. Shively, S. B. et al. Developmentally Arrested Basket/Stellate Cells in Postnatal Human Brain as Potential Tumor Cells of Origin for Cerebellar Hemangioblastoma in von Hippel-Lindau Patients. Journal of Neuropathology and Experimental Neurology 81, 885–899 (2022).

75. Vergauwen, E., Forsyth, R., Vortmeyer, A. & Gläsker, S. Expression of Hemangioblast Proteins in von Hippel-Lindau Disease Related Tumors. Cancers 15, 1–13 (2023).

76. Monzon, F. A. et al. Chromosome 14q loss defines a molecular subtype of clear-cell renal cell carcinoma associated with poor prognosis. Modern Pathology 24, 1470–1479 (2011).

77. Shen, C. et al. Genetic and Functional Studies Implicate HIF1α as a 14q Kidney Cancer Suppressor Gene. Cancer Discov 1, 222–235 (2011).

78. Meléndez-Rodríguez, F. et al. HIF1α Suppresses Tumor Cell Proliferation through Inhibition of Aspartate Biosynthesis. Cell Reports 26, 2257–2265.e4 (2019).

79. Goswami, A., Surve, A. & Venkatesh, P. Optical Coherence Tomography Angiography of Early Stage 1a Retinal Hemangioblastoma in Von-Hippel-Lindau. J Kidney Cancer VHL 8, 15–18 (2021).

80. Mouchtouris, N. et al. Biology of cerebral arteriovenous malformations with a focus on inflammation. Journal of Cerebral Blood Flow and Metabolism 35, 167–175 (2015).

81. Wang, Y. et al. Ephrin-B2 controls VEGF-induced angiogenesis and lymphangiogenesis. Nature 465, 483–486 (2010).

82. G. Hirrlinger, P., Scheller, A., Braun, C., Hirrlinger, J. & Kirchhoff, F. Temporal Control of Gene Recombination in Astrocytes by Transgenic Expression of the Tamoxifen-Inducible DNA Recombinase Variant CreERT2. GLIA 54, 11–20 (2006).

83. Fernández-Chacón, M. et al. iSuRe-Cre is a genetic tool to reliably induce and report Cre-dependent genetic modifications. Nature Communications 10, 1–13 (2019).

84. Acute postnatal ablation of Hif-2α results in anemia. https://www.pnas.org/doi/10.1073/pnas.0608382104 doi:10.1073/pnas.0608382104.

85. Ryan, H. E. et al. Hypoxia-inducible Factor-1α Is a Positive Factor in Solid Tumor Growth. Cancer Research 60, 4010–4015 (2000).

